# PT-seq: A method for metagenomic analysis of phosphorothioate epigenetics in complex microbial communities

**DOI:** 10.1101/2024.06.03.597111

**Authors:** Yifeng Yuan, Michael S. DeMott, Shane R. Byrne, the Global Microbiome Conservancy, Peter C. Dedon

## Abstract

Among dozens of known epigenetic marks, naturally occurring phosphorothioate (PT) DNA modifications are unique in replacing a non-bridging phosphate oxygen with redox-active sulfur and function in prokaryotic restriction-modification and transcriptional regulation. Interest in PTs has grown due to the widespread distribution of the *dnd, ssp*, and *brx* genes among bacteria and archaea, as well as the discovery of PTs in 5-10% of gut microbes. Efforts to map PTs in complex microbiomes using existing next-generation and direct sequencing technologies have failed due to poor sensitivity. Here we developed PT-seq as a high-sensitivity method to quantitatively map PTs across genomes and metagenomically identify PT-containing microbes in complex genomic mixtures. Like other methods for mapping PTs in individual genomes, PT-seq exploits targeted DNA strand cleavage at PTs by iodine, followed by sequencing library construction using ligation or template switching approaches. However, PT-specific sequencing reads are dramatically increased by adding steps to heat denature the DNA, block pre-existing 3’-ends, fragment DNA after T-tailing, and enrich iodine-induced breaks using biotin-labeling and streptavidin beads capture. Iterative optimization of the sensitivity and specificity of PT-seq is demonstrated with individual bacteria and human fecal DNA.

## INTRODUCTION

The prokaryotic epigenome comprises dozens of different pre- and post-synthetic DNA modifications that function in host defense in restriction-modification (RM) systems, in immunity against RM in bacteriophage, and in regulating gene expression (1,2). Phosphorothioate (PT) DNA modifications are unique in replacing a non-bonding phosphate oxygen in the DNA backbone with an oxidizable and nucleophilic sulfur (3–19). It has been more than 50 years since PT modification was first created synthetically as a way to engineer nuclease resistance into nucleic acids (3) and more than 15 years since PTs were discovered to occur naturally in bacterial genomes (18) and more recently in archaea (20). In addition to RM, there is evidence that PTs regulate gene expression, with PT oxidation proposed to provide epigenetic regulation of transcription of redox homeostasis genes (8). PTs also appear to protect bacteria against reactive oxygen and nitrogen species, such as peroxides and peroxynitrite (**Fig. 1D**) (6,8,21) but render the bacteria 5-fold more sensitive to neutrophil-derived hypochlorous acid (HOCl) (**Fig. 1D**) (7). Recent studies have revealed that PT-containing bacteria are present in the gut microbiome (11,15), with mass spectrometric detection of PTs in fecal DNA as PT-linked nucleotides showing temporal variation (15). These observations raise questions about the fate of microbes bearing an abundant and chemically-reactive PT epigenetic mark in both the healthy gut and in the chronic inflammation of inflammatory bowel disease (IBD) and other pathologies (12,22,23).

**Fig. 1.**
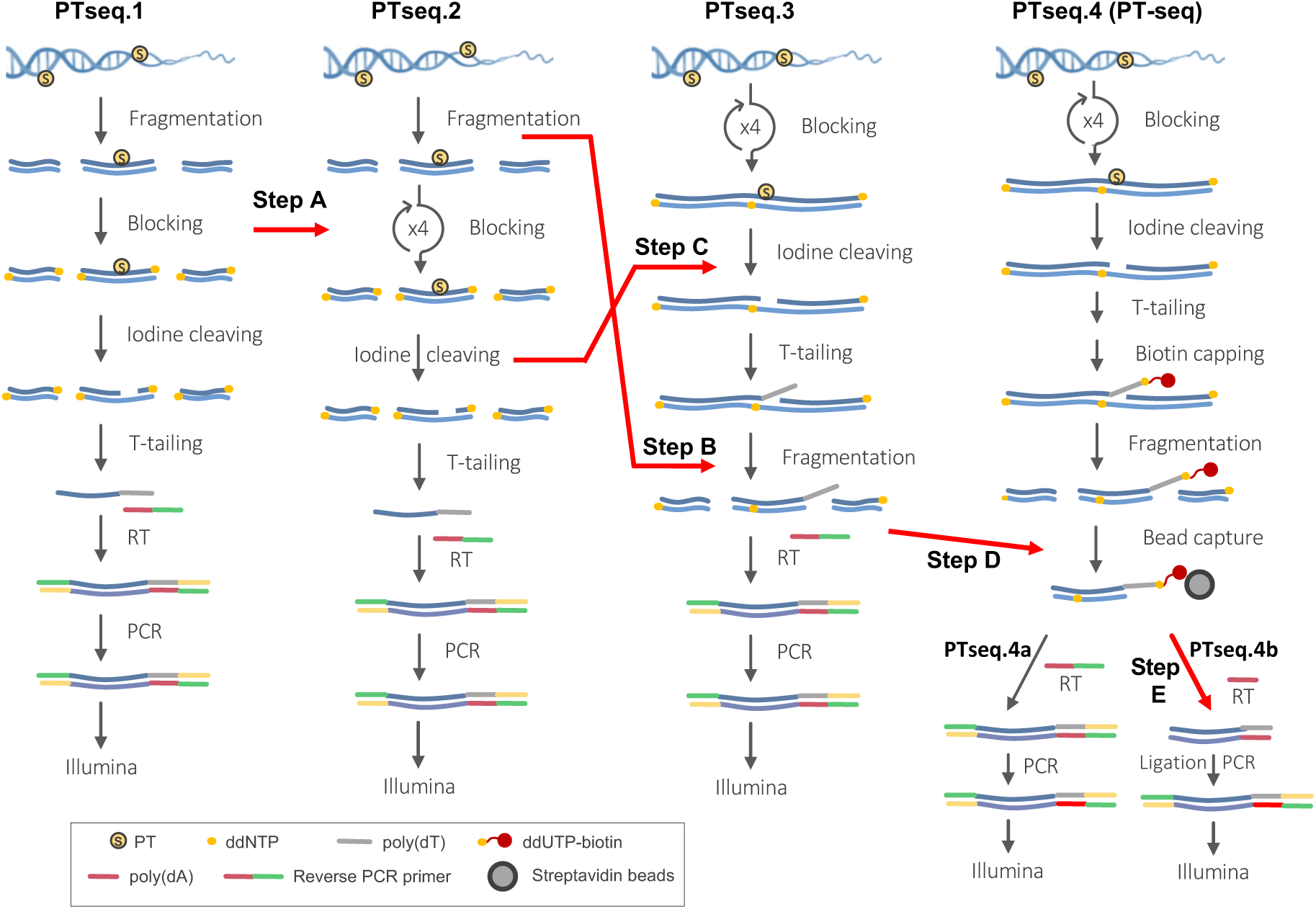
Iterative optimization of PT-seq. The original T-tailing component of Nick-seq was iteratively optimized for DNA fragmentation, iodine cleavage conditions, and affinity purification of T-tailed DNA in 3 discrete protocols (PTseq.1-Ptseq.4). The optimized steps are indicated with red arrows.

To attempt to answer these questions, a variety of methods have been developed for mapping PTs at single nucleotide resolution in prokaryotic genomes. These methods involve either next-generation sequencing (NGS) of chemically-derivatized DNA or single-molecule real-time (SMRT) sequencing of native PTs (5) (**Table 1**). The original NGS method, termed iodine-coupled deep sequencing (ICDS), used iodine to site-specifically oxidize PTs to produce DNA double-strand breaks at sites with PTs on both strands, as in the consensus sequence G_PS_AAC/G_PS_TTC in *Escherichia coli* B7A (5). The double-strand breaks were then blunt-end ligated to double-strand linkers for PCR amplification and NGS sequencing (5). A variant of ICDS without the enrichment process, named PT-IC-seq, could quantitatively determine the PT modification percentage (9). Unfortunately, these methods only worked for microorganisms where modifications simultaneously occurred on opposing strands and did not work for microorganisms in which PTs occur on only one strand of the consensus sequence, such as C_PS_CA in *Vibrio cyclitrophicus* FF75 (5). To capture both single- and double-stranded PTs, we developed Nick-seq (24), which involved blocking of pre-existing strand-breaks and iodine oxidation of PTs, followed by both 3-extension by nick-translation with PT-modified dNTPs to produce nuclease-resistant oligonucleotides and 3’-terminal transferase tailing with a polymer of dTMPs in a template independent manner. Following library preparation for both samples and NGS, the complementary datasets were mined with a custom workflow to increase sensitivity, specificity, and accuracy of the map (24). However, Nick-seq proved to be too insensitive to capture PT-containing genomes comprising 5-10% of the gut microbiome (24).

**Table 1.**
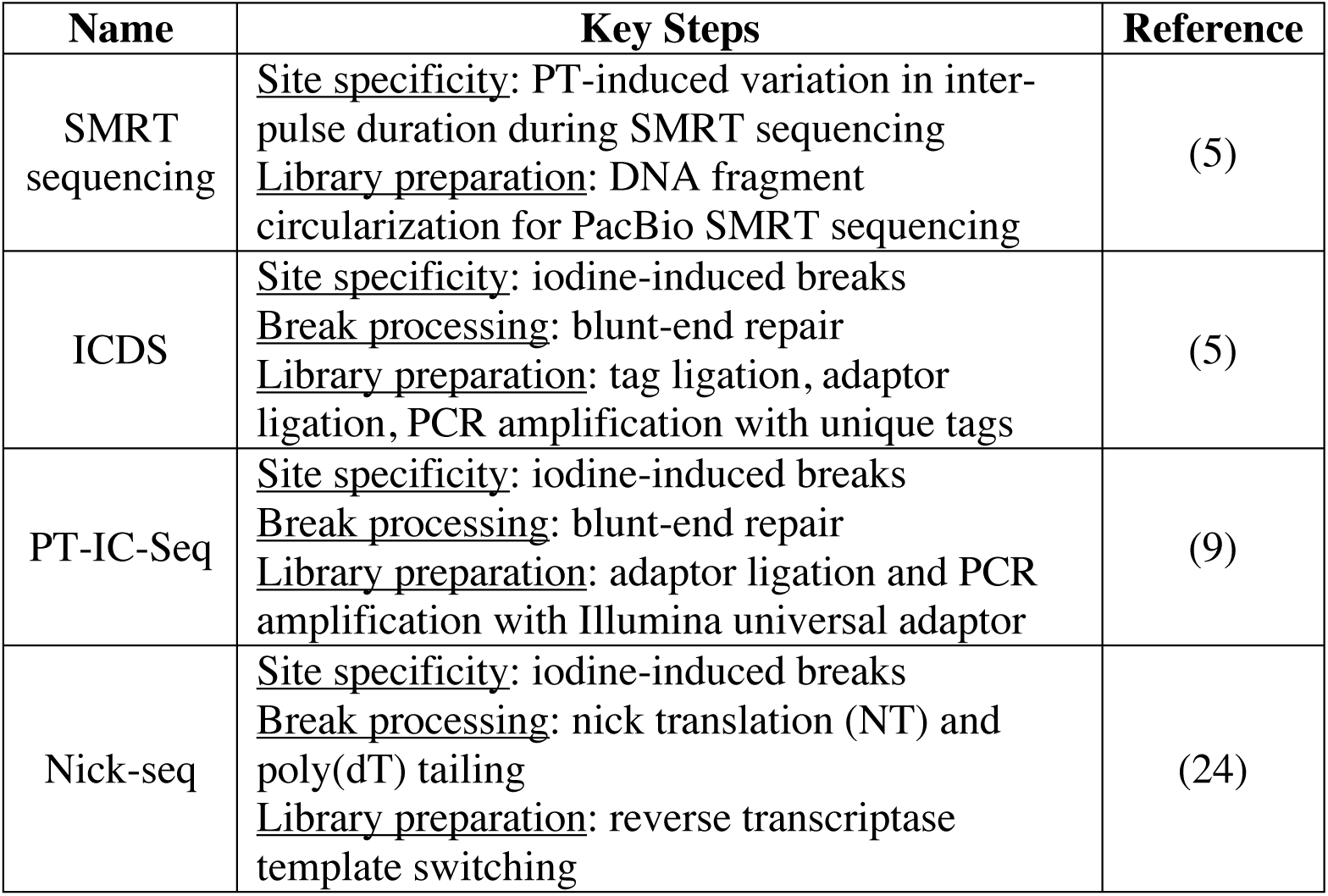
Single-nucleotide resolution PT sequencing methods.

Given the problems with these PT sequencing methods, we set out to optimize critical steps in the T-tailing portion of Nick-seq to maximize specificity and sensitivity. Here we present this development process in four iterations that demonstrate the critical features of the method and progressively improve the sensitivity and specificity of the method by orders-of-magnitude.

## RESULTS

### The PT-seq protocol

Here we present the final, optimized PT-seq protocol (PTseq.4a) derived from the four iterative adjustments depicted in **Figure 1**. The point is to describe the optimized PT-seq as a context for understanding the impact of changes that occurred in the three preceding optimization steps.

### Block pre-existing strand breaks

1) DNA (5-10 µg) is diluted to 30.5 µl in filtered H_2_O and subjected to four blocking cycles.

2) One blocking cycle consists of the following:

a. DNA is heated at 94 °C for 2 min and immediately cooled on ice for exactly 2 min.

b. rSAP (1 µl) and rCutSmart buffer (3.5 µl) are added, the reaction is incubated for 30 min at 37 °C, and the phosphatase is then heat deactivated for 10 min at 65 °C

c. Four ddNTPs (1 µl, 2 mM each), TdT reaction buffer (1.5 µl), CoCl_2_ (5 µl, 0.25 mM), terminal transferase (1 µl, 20 units), and filtered H2O (3.5 µl) are added, and the reaction incubated for 60 min at 37 °C.

[Note: fresh reagents are added to each subsequent cycle at specific steps: rSAP (1 µl) and rCutSmart buffer (0.3 µl) were added at step b, while ddNTPs (0.3 µl each), Tdt reaction buffer (0.3 µl), CoCl_2_ (0.3 µl), and terminal transferase (1 µl) were added at step c.]

3) After all cycles are complete, excess unincorporated ddNTPs are removed by treatment with a DNA Clean & Concentrator kit (Zymo #11-304C).

### Iodine cleavage

4) Blocked DNA (32 µl) is incubated with 500 mM Tris-HCl pH 9.0 (4 µl) and iodine solution (4 µl, 5 mM) for 10 min at 25 °C and placed on ice.

5) The reaction is split into two 20 µl aliquots and each is filtered using a single DyeEX column to remove salts and iodine, followed by recombining eluents.

### dT-tailing

6) The DNA is denatured by heating at 94 °C for 2 min and then cooling on ice for exactly 2 min.

7) The reaction is adjusted with rCutSmart buffer (5 µl), rSAP (1 µl), and filtered H_2_O to a final volume of 50 µl, and incubated for 30 min at 37 °C to remove 3ʹ-terminal phosphates arising from iodine cleavage.

8) The phosphatase enzyme is then deactivated for 10 min at 65 °C.

9) The reaction is adjusted with dTTP (1 µl, 1 mM), TdT reaction buffer (1 µl), CoCl_2_ (6 µl, 0.25 mM), filtered H_2_O (1 µl), and terminal transferase (1 µl, 20 units) and incubated for 45 min at 37 °C to add dT-tails to the 3’-terminal ends created specifically by iodine at PT sites.

10) The reaction is divided into three 20 µl aliquots, excess dTTP is removed using three DyeEX columns, and then eluents are recombined.

### ddUTP-biotin conjugation

11) The reaction (60 µl) is adjusted with TdT reaction buffer (7.8 µl), CoCl_2_ (7.8 µl), ddUTP-biotin (1 µl, 1 mM), filtered H_2_O (0.4 µl), and terminal transferase (1 µl) and incubated for 60 min at 37 °C to terminate the dT-tails with a conjugated biotin moiety.

12) The DNA sample is divided into 4 aliquots and each is cleaned with a separate DyeEX column. Combine all four eluents.

### DNA fragmentation

13) The processed DNA is diluted in filtered H_2_O (500 µl) in a 2ml Eppendorf tube and kept in an ice-bath at all times to dissipate heat during sonication.

14) DNA is fragmented by probe sonication (Branson Sonifier 250 Ultrasonic Cell Disruptor outfitted with a microtip probe; 50% duty-cycle and an output control setting to attain ∼20-25 on the meter) for 5 cycles (1 cycle = 2 min ON followed by 1 min OFF).

15) The sample is dried using a SpeedVac (∼2.5 h at ∼450 mTorr)

### Streptavidin bead capture

16) DNA fragments are mixed with Hydrophilic Streptavidin Magnetic Beads (5 µl) and binding buffer (500 µl) (5 mM Tris-HCl, pH 7.5, 1 M NaCl, 0.5 mM EDTA)

17) The sample is incubated for 60 min on a Nutator at ambient temperature.

18) The beads are subjected to a pull-down process using a magnetic separation rack and washed three times with binding buffer (100 µl).

19) After discarding the supernatant, beads are resuspended in filtered H_2_O (20 µl).

### Illumina library preparation

20) Finally, the enriched DNA is directly subjected to Illumina library preparation using a SMART ChIP-seq kit (Takara, #634865) following the manufacturer’s protocol.

21) The final PCR step is performed using Illumina primers provided in the kit, and 12 cycles are typically run for amplification.

22) The sample is submitted for paired-end sequencing (150-bp), using a Custom Read2 sequencing primer (poly(dA) followed by universal read 2 primer) that was supplied in the kit.

The following sections detail the optimization and refinement steps in developing this PT-seq protocol.

### Blocking pre-existing 3’-OH termini

The first optimization step addressed the efficiency of blocking 3’-ends of pre-existing strand breaks with dideoxy-NTPs and terminal transferase (**Fig. 1, PTseq.1**). Left unblocked, the 3’-OH ends will behave identically to cleaved PT sites and contribute to the background signal in downstream Illumina sequencing results. In most published methods, a single blocking cycle is considered satisfactory. However, we hypothesized that repeated heat denaturation and blocking would uncover and block additional 3’-ends more efficiently given possible sequence-specificity of terminal transferase as observed with ligases (25). To test this idea, we developed an assay in which the blocking efficiency was calculated by measuring incorporated ddNMPs (**Fig. S1B**). The workflow started with 1 to 7 cycles of heat denaturation, cooling, and blocking with a mixture of ddATP, ddCTP and ddTTP and terminal transferase. After removal of excess ddNTPs, DNA was again capped with only ddGTP to act as a final mark. Finaly, after another round of cleaning, the incorporated ddNMPs were quantified by chromatography-coupled triple quadrupole mass spectrometry (LC-MS/MS) (26). Clearly, based on a decreasing quantity of ddGMP being incorporated after cycle 1 (**Fig. S1B**), one round of blocking was deemed insufficient to maximally remove pre-existing breaks. Further cycles showed a continuing decrease of incorporated ddGMP through cycle 3 (**Fig. S1B**), after which we observed a slight increase in the total incorporation of all ddNMPs before peaking. Subsequently, the new PTseq.2 protocol adopted 4 blocking cycles using all four ddNTPs (**Fig. 1, Step A**).

### Optimizing DNA fragmentation

NGS read depth is optimal with a DNA fragment range of ∼200-500 bp, which can be achieved by enzymatic or mechanical means. Sonication-based fragmentation was used in Nick-seq and other PT sequencing methods (5,9,24). However, previous studies revealed that PT was extremely sensitive to oxidation (4,7). Since ultrasonication is known to generate reactive oxygen species (27), we hypothesized that DNA fragmentation by probe sonication would consume PTs. To test this idea, we defined a 10-min probe sonication condition (settings detailed in Methods) that produced DNA fragments of ∼200-500bp (**Fig. S2B**). We then compared PT levels under these conditions to probe sonication for 0 and 2 min. As shown in **Figure S2C**, the 10 min sonication significantly lowered the abundance of PT dinucleotides.

Since probe sonication is the most effective means to produce NGS-friendly DNA fragmentation, we decided to move the sonication step in PTseq.2 after iodine cleavage and subsequent end labeling at cleaved PT sites in a new protocol PTseq.3 (**Fig. 1**). Sonication-induced strand breaks would thus not lead to any signal. Next we optimized the iodine-induced PT cleavage step.

### Optimizing iodine reaction conditions

Iodine-based cleavage of DNA at PTs was originally developed in 1988 by Eckstein and coworkers (28) and has been used in all NGS-based PT mapping methods (5,9,24) (**Fig. 1, Step C**). However, conditions of concentration, temperature, time, and sequence-dependence for optimal cleavage efficiency have not been defined, which is important since prior studies with other oxidants suggests that the conversion of PTs to DNA strand breaks is not stoichiometric (7). To assess fragmentation efficiency, we used HPLC to monitor the three major components of a PT cleavage reaction (**Fig. S3**): an uncut oligo containing a single PT as the starting material (e.g., AG*TCA), a desulfurated oligo where the PT is converted to a phosphate (14) (e.g., AGTCA), and release of di- or tri-nucleotides to denote successful cleavage of the oligo (e.g., AG). Using this assay, we found that the minimum concentration of iodine that ensured complete reaction was one I_2_ molecule for each nucleotide in the reaction with the single-stranded AG*TCA oligonucleotide (**Fig. S7A**).

We next addressed the role of PT sequence contexts on iodine cutting efficiency using a set of 5-mer oligonucleotides representing all 16 possible sequence contexts (**Fig. S7B**), with cutting efficiency quantified by release of 3-mer fragments (e.g., CCA, CCC, CCG, and CCT). As shown in **Figure S7B**, cutting efficiencies ranged from 38.9% to 98.4%. These cutting efficiencies with a CCXXA context are higher than those observed in the AGXCA motif (∼25%; **Figs. S7, S8**), which suggests that sequence effects more distant than nearest neighbors play a role in the PT cleavage efficiency.

We also assessed the structure dependence of iodine reactivity. Here we compared single- to double-stranded oligonucleotides using three longer sequences that represented a range of cutting efficiencies noted in **Figure S7B**. Each possessed a melting temperature greater than 27°C when paired with a corresponding complement (**Fig. S7C**), so we performed this experiment at 15°C with excess non-PT complementary strand to ensure double-stranded substrates for the iodine reaction. We compared these results with parallel reactions that lacked the complementary strands. We observed no difference between single- and double-stranded substrates for the sequence contexts tested (**Fig. S7C**).

With the optimal iodine concentration, we next optimized incubation temperature and reaction time. As shown in **Figure S8A**, there was little difference in cleavage efficiency at 0 °C (ice), room temperature, or 65 °C. Although incubation at room temperature produced slightly more PT conversion to phosphate, we adopted it as a matter of convenience. For reaction time, we found little difference between 5 min and overnight incubations (**Fig. S8B**). Again, we confirmed that the iodine-to-nucleotide ratio must be at ≥1 to eliminate all of the starting PT-containing oligonucleotide (**Fig. S8B**). Interestingly, we observed that desulfuration to phosphate increased slowly over time, perhaps due to some kind of intermediate that was difficult to identify and track on the corresponding chromatographic profile. We concluded that the iodine-PT cutting reaction occurred rapidly and to completion within a few minutes, if not faster.

Finally, we optimized the concentration of Tris(hydroxymethyl)aminomethane buffer in the reaction. Tris had been shown to be critical for iodine cleavage of PT by forming a covalent intermediate with the sulfur (Tris-N graft), which rearranges to displace the sulfur, driving the reaction forward to hydrolysis (14). In the absence of Tris, the reaction favors the desulfuration pathway. Indeed, we observed no evidence for AG release in the absence of Tris (**Fig. S8C**). However, with increasing Tris concentration, release of AG continued to improve while the desulfuration product moved in the opposite direction (**Fig. S8C**).

Along with optimized DNA fragmentation (**Fig. 1, Step B**), we added the optimal conditions for the iodine cleavage to the new protocol PTseq.3 (**Fig. 1, Step C**). However, the T-tailing step of PTseq.2 and PTseq.3 proved to be problematic, as we discovered next.

### T-tailing causes false positive PT sites and is inefficient at single-stranded PTs

Over the course of testing the optimized protocols, we discovered two problems with the T-tailing step. The first involved false PT positive sites at endogenous runs of dT. In the template switching/reverse transcription step of PTseq.1-3 (**Fig. 1**), an off-target effect of the poly(dA) primers used to recognize the added poly(dT) tails was that the primers could also anneal to naturally occurring poly(dT) sequences within the genome, leading to PCR amplification of these sites. This was demonstrated by applying PTseq.2 to genomic DNA isolated from *Lachnospiraceae sp.*, which we found by LC-MS/MS to possess PTs at G_PS_A and G_PS_C motifs at 70 and 153 PTs per 10^6^ nt, respectively (16). With a pileup depth cutoff of 10 (see METHODS section below), PTseq.2 detected 89 sites containing G_PS_A and 147 sites containing G_PS_C (per 10^6^ nt) in a *Lachnospiraceae sp.*, which is similar to the LC-MS/MS data (16). However, only 44% of read pileups were detected at the correct 5ʹ-G_PS_AGC/ G_PS_CTC-3ʹ motif, with 42% of read pileups detected at internal NTTTT sites (**Fig. 2A, Supplementary Table S4**). An example of this misalignment at a specific NTTTT site is shown in **Figure S9A**. The dilution of meaningful reads with false-positives at NTTTT sites will greatly reduce sensitivity of PT-seq for complex mixtures of genomes.

**Fig. 2.**
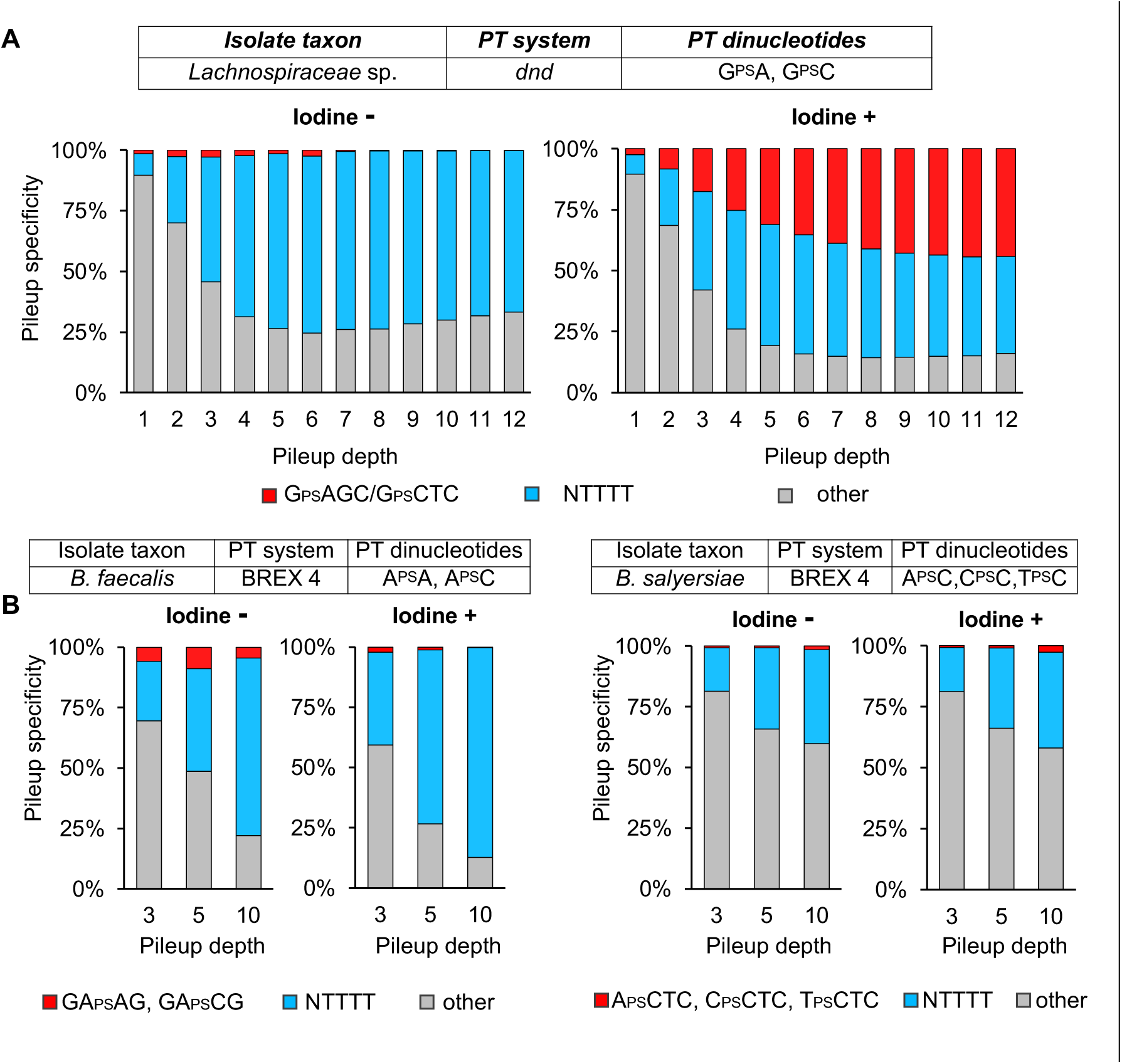
PTseq.2 displayed limited specificity in gut microbiome isolates containing single-stranded PT motifs. Three PT-containing gut microbiome bacterial isolates were sequenced using Ptseq.2. The PT gene family and PT dinucleotides identified by LC-MS/MS are noted for each isolate. (**A**) Less than 50% of read pileups were PT-specific (red) in *Lachnospiraceae* sp.at G_PS_AGC and G_PS_CTC. (**B,C**) Less than 5% of read pileups were PT-specific (red) in *Butyricimonas faecalis* (**B**) and *Bacteroides salyersiae* (**C**). In all cases, an aliquot of sample without iodine treatment (Iodine -) was used as control to show PT-specific read pileups. Endogenous NTTTT sites were the primary source of non-specific reads (gray).

Another problem noted with T-tailing was the inefficiency of the template-independent polymerase, terminal transferase (TdT). We noted this in the application of PTseq.2 to genomic DNA from two PT-containing isolates, *Bacteroides salyersiae* DSM18765 and a putative *Butyricimonas faecalis* strain, both of which harbor the BREX type 4 PT modification system (16). The read pileups at conserved PT motifs accounted for less than 5% of all reads (**Fig. 2B**), suggesting inefficient dT-tailing. As illustrated by Motea (29), the structure of TdT forms a “thumb-finger-palm” shape (**Fig. S9B**), with the finger and palm domains forming a “lariat-like” loop limiting the interaction with duplex DNA. This structure suggests that dNTPs will be added more efficiently to 3’-protruding ends than to 3’-recessed ends, to blunt ends, or especially to single-stranded nicks (**Fig. S9C**). In support of this model, the low (5%) read pileup observed for *B. salyersiae* and *B. faecalis* may result from the fact that the PT sites all have single-stranded modifications (GA_PS_AG, GA_PS_CG, A_PS_CTC, C_PS_CTC, and T_PS_CTC) and the iodine-induced breaks would be inefficiently tailed by TdT. In addition, the low sensitivity and specificity of the PTseq.2 protocol for *Lachnospiraceae sp.* mentioned earlier correlates with the fact that only a small proportion of PTs occurred at double-stranded PT motifs (5ʹ-G_PS_AGC/ G_PS_CTC-3ʹ) and even these would produce only slightly more accessible 3’-recessed ends (**Fig. S9D**). We next explored an approach to reducing read pileup noise and increasing the intensity of meaningful reads.

### Enrichment of T-tailed PT cleavage sites

The false positive read pileups caused by genomic T runs and the inefficiency of TdT for single-strand breaks at PTs led us to add a step to enrich for poly(T) tails at PT break sites and thus increase the signal-to-noise ratio in the sequencing read pileups. Here we modified PTseq.3 by using a biotin-conjugated ddUMP to terminate elongation of the dT-tails (**Fig. 1, Step D**). Following sonication-induced DNA fragmentation, the biotin-labeled DNA fragments are then captured on streptavidin coated magnetic beads and the bulk of the unlabeled DNA fragments are washed away. The purified T-tailed DNA is then subjected to template switching reverse transcription and PCR amplification for sequencing. This affinity purification step reduces contributions from genomic T runs and enriches for meaningfully T-tailed fragments representing PT sites. The combined set of optimizations to this point is now referred to as PTseq.4a in **Figure 1D**; the relevance of PTseq.4b is addressed shortly.

The effect of biotin capture enrichment was apparent in applications of PTseq.4a to the *B. salyersiae* genomic DNA studied earlier. PTseq.4a identified 3 motifs (A_PS_CTC, C_PS_CTC and T_PS_CTC) corresponding to PT dinucleotides A_PS_C, C_PS_C and T_PS_C detected by LC-MS/MS analysis (16). With a pileup depth cutoff of 10 to define a PT site, a total of 1636 PTs (per 10^6^ nt per 10^7^ reads) were detected by PTseq.4a compared to 3 PTs with PTseq.2 (**Supplementary Table S4**). At the same time, the specificity of pileups at conserved motifs increased from 2.7% with PTseq.2 to 95% with PTseq.4a (**Fig. 3A,C; Supplementary Table S4**).

**Fig. 3.**
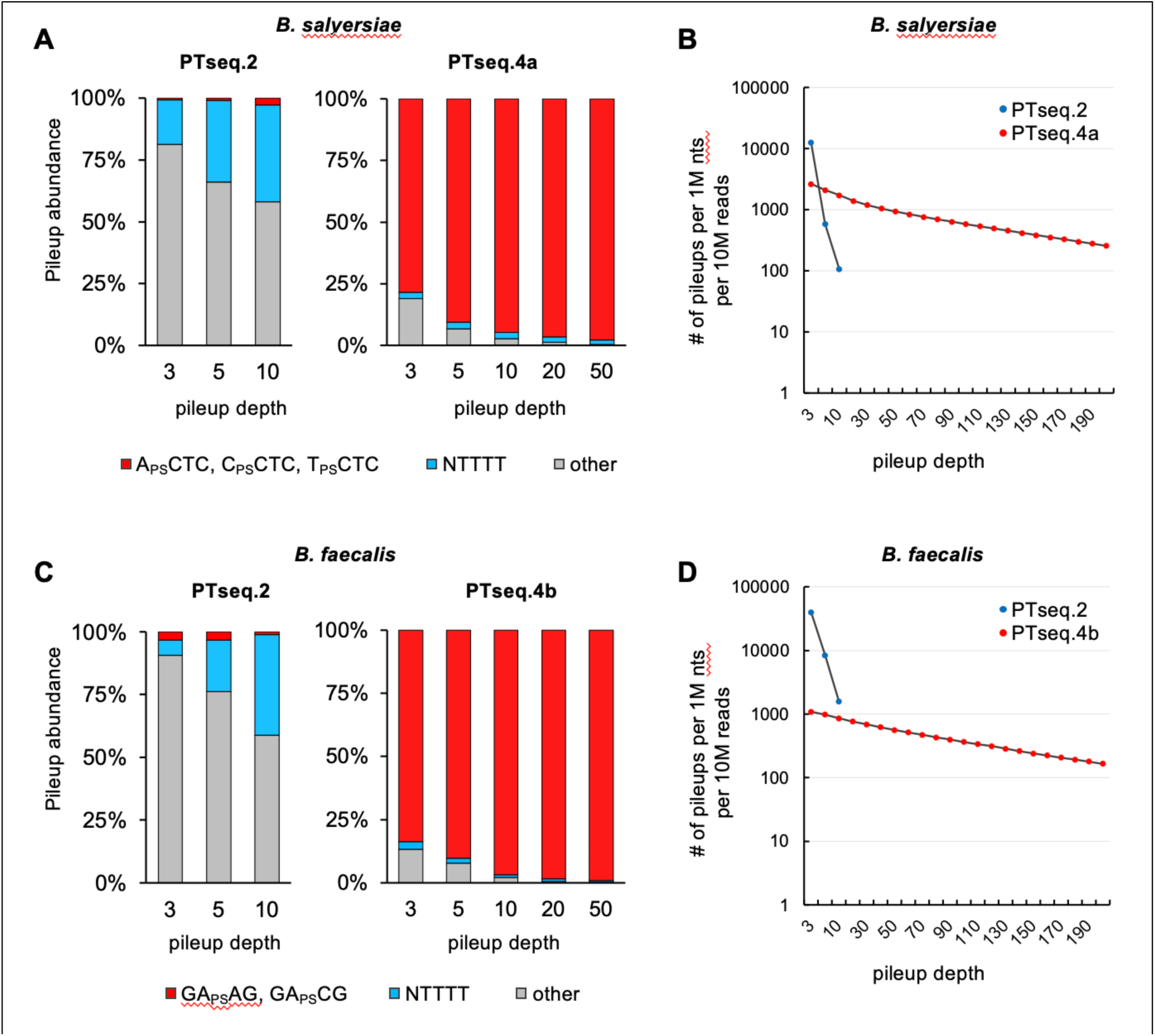
PTseq.4 displayed high sensitivity and specificity in bacterial isolates, but Ptseq.4a outperformed Ptseq.4b. **(A,C)** >90% of read pileups involved PT-specific sites (red) in genomic DNA from in *B. salyersiae* (**A**) and *B. faecalis* (**C**) when using PTseq.4a and PTseq.4b, respectively, compared to < 5% specificity for PTseq.2. (**B,D**) Read pileups as a function of pileup depth were dramatically higher for Ptseq.4 compared to Ptseq.2 for both *B. salyersiae* (**B**) and *B. faecalis* (**D**), with Ptseq.2 limited to a maximum pileup of 20-25 reads. PTseq4a outperformed PTseq.4b by 2- to 3-fold.

### Optimized processing of affinity-purified T-tailed DNA fragments

Finally, there are several options for processing the affinity-purified, T-tailed DNA fragments for sequencing, here on the Illumina Novaseq platform (**Fig. S4**). One option used in the PTseq.1-3 and PTseq.4a protocols discussed earlier is the template-switching/reverse transcriptase SMART ChIP-seq kit (Takara, #634865; **Fig. S4** left). This kit is less cost effective for small numbers of samples than the NEBNext Ultra II DNA library prep kit (**Fig. S4** right). Here we tested the performance of NEBNext Ultra II DNA library prep kit using PTseq.4 with genomic DNA from *B. salyersiae* and *B. faecalis* and then using the kit to synthesize the first cDNA strand by reverse transcription directly on the streptavidin beads after capture and washing. This added refinement, now referred to as PTseq.4b (**Fig. 1**), identified two major motifs GA_PS_AG and GA_PS_CG that corresponded to PT dinucleotides A_PS_A and A_PS_C identified by LC-MS/MS analysis (16). It also revealed a trace amount of CT_PS_TC and CG_PS_TC of which the corresponding PT dinucleotides were below the detection limit of LC-MS/MS analysis (**Supplementary Table S4**). With a pileup depth cutoff of 10, a total of 834 PT sites (per 10^6^ nt per 10^7^ reads) were detected using PTseq.4b, compared with that of 4 PT sites when using PTseq.2 (**Supplementary Table S4**). At the same time, the specificity of pileups at conserved motifs increased from 1.1% with PTseq.2 to 97% with PTseq.4b (**Fig. 3C, Supplementary Table S4**). However, PTseq.4a slightly outperformed PTseq.4b by a factor of 2-3 in terms of read pileups (**Fig. 3B,D**), so we opted to use PTseq.4a for subsequent work.

### PTseq.4a exhibits optimal sensitivity and specificity for PT mapping in the human gut microbiome

To assess the benefits of the various refinements and optimizations of the PT sequencing protocol, we applied all of the PTseq protocols in **Figure 1** in parallel to a fecal DNA sample obtained from a healthy human donor and analyzed by LC-MS/MS for PT dinucleotides (15). PTseq.4a identified 16 sites of C_PS_AG, 22 sites of C_PS_CA, 0.02 sites of G_PS_AAC, 3.7 sites of G_PS_AGC, 9.1 sites of G_PS_ATC, 2.8 sites of G_PS_CTC, and 0.01 sites of G_PS_TTC per 10^6^ nt in 26 genomes (**Supplementary Table S5**) using a pileup depth cutoff of 5 (**Fig. S10A, Supplementary Table S6**). This agrees well with results from LC-MS/MS analysis: 8 C_PS_A, 22 C_PS_C, and 12 G_PS_A per 10^6^ nt (**Fig. S10B, Supplementary Table S7**). The discrepancy with C_PS_A between PTseq.4a and LC-MS/MS could be explained by misidentification of some pileups at C_PS_CAG sites instead of CC_PS_AG sites, given that they are identical motifs with modification sites only one residue apart. Such discrepancies need careful consideration during data mining. In addition, we were unable to detect G_PS_C or G_PS_T during LC-MS/MS analysis, but since the frequencies of these sites were low in PTseq.4a, they were likely below the lower limit of detection for LC-MS/MS.

Finally, the PTseq.1-3 methods (**Fig. 1**) identified fewer PT-containing genomes (**Supplementary Table S5**) and exhibited more limited sensitivity and specificity compared with PTseq.4a (**Fig. S10**). After normalizing by read numbers, PTseq.4a demonstrated the best sensitivity and specificity (**Fig. 4**), with PTseq.4a achieving a >200-fold increase in sensitivity and a 35-fold increase in specificity in microorganisms that contained mostly single-stranded PT motifs compared with PTseq.2 (**Fig. 4C,D**). For example, 1.7, 2.7, 9.7, and 104.1 PT sites were identified (per 10^6^ nt per 10^7^ reads) using PTseq.1, PTseq.2, PTseq.3, and PTseq.4a, respectively, with a pileup depth cutoff of 5 (**Fig. 4B**). The corresponding specificity of pileups at conserved motif sites was 19.2%, 25.3%, 35.6%, and 47.2%, respectively (**Fig. 4, Supplementary Table S6**). Moreover, the fold-change increase for motif sites containing C_PS_A and C_PS_C was much higher when using PTseq.4a compared with the other methods, suggesting that PTseq.4 was able to identify single-stranded C_PS_AG and C_PS_CA motifs more efficiently.

**Fig. 4.**
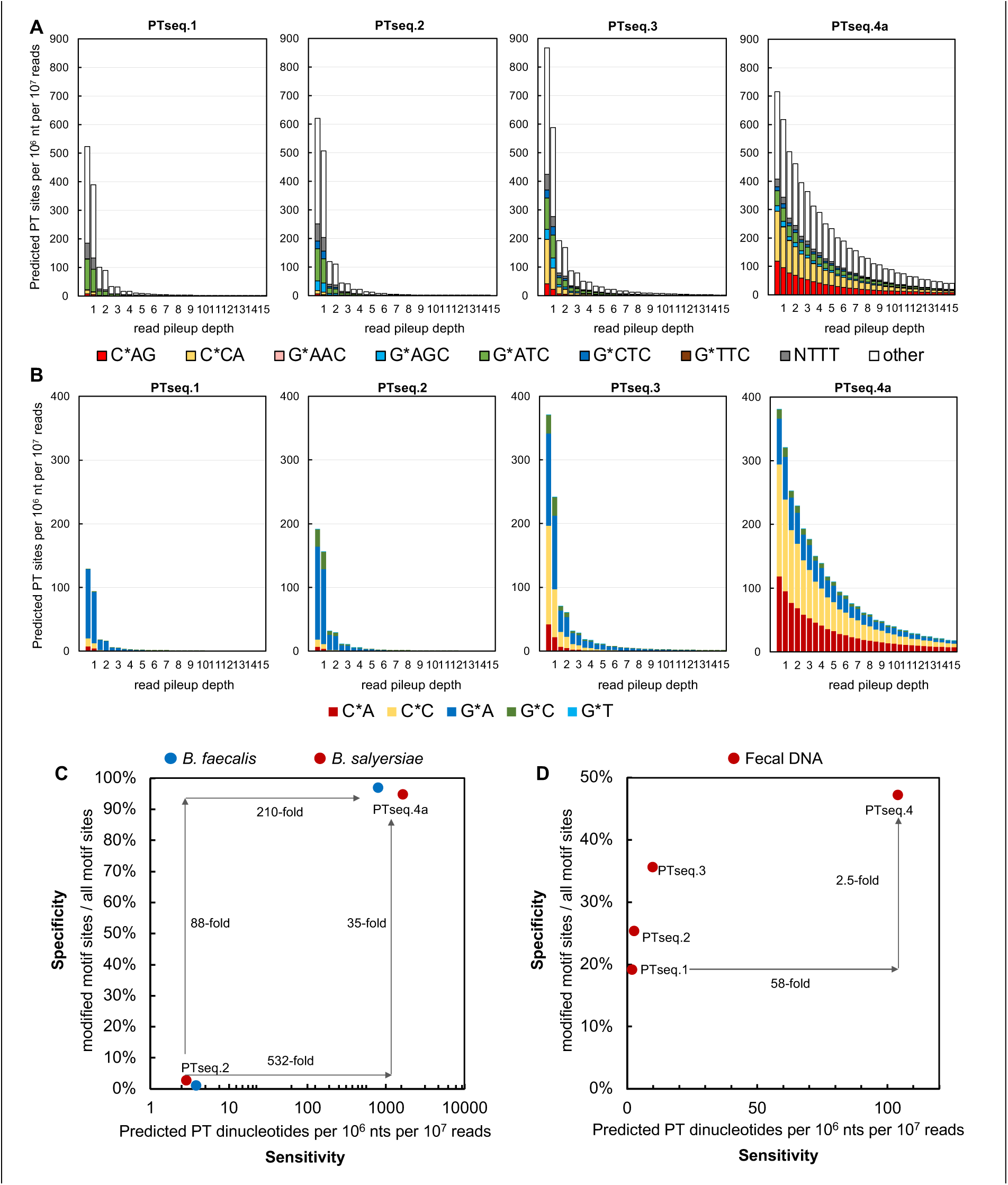
Comparison of PT-seq protocol performance with a human gut microbiome sample. (**A,B**) Graphs depict the number of PT-modified conserved motifs (A) and PT dinucleotides (B) per 10^6^ nt per 10^7^ reads decreased as a function of cutoff of read pileup depth. The sensitivity (number of read pileups) and specificity (accuracy of calling PT motifs) increased progressively from Ptseq.1 to Ptseq.4. (**C,D**) PTseq.4 surpassed the other protocols in sensitivity and specificity in both bacterial isolates (**C**) and a human fecal DNA sample (**D**). The number of PT dinucleotides and motif sites were normalized for comparison (per 10^6^ nt per 10^7^ reads).

## DISCUSSION

In summary, we performed an exhaustive step-by-step optimization of the T-tailing method of our Nick-seq protocol to improve sensitivity and specificity of mapping PT sites in complex mixtures of genomes, such as the gut microbiome. These refinements included (1) four blocking cycles to lower background signals from spurious PT sites, (2) optimized conditions for iodine cleavage, (3) DNA fragmentation by sonication after dT-tailing to avoid oxidation and thus a reduction of actual PT sites, and (4) enriching for T-tailed DNA fragments using biotin-streptavidin affinity purification (**Fig. 1, Steps A-D**).

The goal of these optimization efforts was to achieve a method that would allow meaningful metagenomic identification of PT-containing bacteria and genome-wide mapping of the PTs in a complex system like the human gut microbiome. We know from LC-MS/MS analysis of fecal DNA (16) that PT-containing bacteria represent 5-10% of the hundreds of different species in the mouse and human gut microbiomes (30,31), which immediately dilutes the PT-containing fragments in fecal DNA by 10- to 20-fold. This dilution is further increased when the variety of bacterial strains is considered among the trillions of individual microbes in the gut microbiome (32). The point here is the need for sensitivity and thus meaningful read pileups in PT-seq sequencing datasets. We showed that PTseq.4a could achieve an approximately 58-fold increase in sensitivity and a 2.5-fold increase in specificity compared with the original method (**Fig. 4**). Ultimately, we concluded that PTseq.4a, now termed PT-seq, exhibited the best sensitivity and specificity, especially for bacterial isolates that harbored predominantly single-stranded PT motifs for which iodine cleavage generates single-strand breaks.

Of course, PT-seq is not without drawbacks that limit the quantitative interpretation of the metagenomic taxonomy of PT-containing bacteria. We showed that there are 4-fold sequence-dependent differences in iodine-induced PT cleavage and variable efficiency of TdT with different cleavage site structures. If these effects hold up in long DNA fragments, then some sequence contexts will be underrepresented in PT-seq read pileups, which limits absolute quantification of the PT-containing strains. This can be partially corrected by weighting according to the sequence-dependent reactivity (e.g., **Figure S7B)** and by comparing the metagenomic data to the absolute quantification of PT dinucleotides by LC-MS/MS. These experimental variances can also be compensated by considering relative changes in PT-bearing bacterial strains among people and during the temporal variation in PT dinucleotide levels we observed in several healthy donors (16).

Over the course of testing these refinements, we also discovered an alternative way to potentially identify PT-containing microbes in the human gut microbiome. We observed that 16S rRNA genes were often hypermodified with PTs compared to other features of genomic architecture (16). Thus, treatment with iodine would target and potentially eliminate 16S rRNA genes located specifically in PT microorganisms (**Fig. S6**). Comparisons of the abundance of OTUs between mock control and iodine treated samples of human fecal DNA revealed a decrease in abundance for *Alistipes putredinis*, a few *Lachnospiraceae* sp., *Bacteroides caccae*, *Ruminococcaceae*, and others (Supplementary Table 8). Interestingly, these results were consistent with PT-containing OTUs that were identified by PT-seq (i.e., PTseq.4a; **Supplementary Table S5**). This approach is a more cost-efficient way to provide a general profile of PT-containing OTUs.

## MATERIAL AND METHODS

### Strains and fecal samples

Cell pellets of human gut microbiome isolates in the Global Microbiome Conservancy (GMbC) collection (33), *Lachnospiraceae sp.* and *Butyricimonas faecalis*, were provided by OpenBiome (openbiome.org, Somerville, MA). Cell pellets of *Bacteroides salyersiae* DSM 18765 were generously provided by Dr. Laurie Comstock (University of Chicago, Chicago, Il.). Human fecal DNA was obtained as part of a parallel study in the PI’s laboratory as previously described (15).

### Extraction of bacterial DNA from bacterial cell culture

Bacterial gDNA was extracted using the E.Z.N.A. Bacterial DNA Kit (#D3350-02) purchased from Omega Bio-tek, Inc. (Norcross, GE), according to the manufacturer’s protocol. RNA was degraded with 2.5 µl of RNase A (Invitrogen #12091021).

### Extraction of bacterial DNA from human fecal samples

An optimized extraction protocol was utilized as previously described (15). Fecal material (180-220 mg) was combined with 0.1 mm (0.3 g) and 0.5 mm (0.3 g) Zirconia beads and 1 mL of InhibitEx buffer from the QIAmp Fast DNA Stool Mini Kit (QIAGEN #51604) in a 2 mL tube. The mixture was processed on a FastPrep Cell Disruptor for 45 seconds using a speed setting of 6.0 (2 total cycles of bead beating). The mixture was then incubated at 95 °C for 15 min. The mixture was again homogenized on a FastPrep Cell Disruptor for 45 seconds using a speed setting of 6.0 (2 total cycles of bead beating). Solid material was pelleted *via* centrifugation at 4 °C for 5 min at 16,100 g. The supernatant was transferred to a new 2 mL tube and kept on ice. InhibitEx buffer (300 µl) was added to the pellet and the mixture was again homogenized on a FastPrep Cell Disruptor for 45 seconds using a speed setting of 6.0 (2 total cycles of bead beating). Solid material was pelleted via centrifugation at 4 °C for 5 min at 16,100 g. The supernatant was pooled with the previous supernatant. 260 µl of 7.5 M ammonium acetate (Sigma-Aldrich #A2706) was added to the pooled supernatants, and the solution was vortexed and incubated on ice for 5 min. Precipitate was pelleted by centrifugation at 4 °C for 10 min at 16,100 g. The supernatant was removed and aliquoted into separate 2 mL tubes (650 µl each). One volume (650 µl) of isopropanol was added to each aliquot, the mixtures were vortexed, and incubated on ice for 30 min. DNA was pelleted by centrifugation at 4 °C for 15 min at 16,100 g. The supernatant was discarded, and pellets were gently washed with 70% ethanol (500 µl) by inverting 3 times before centrifugation at 16,100 g at 4 °C for 15 min. The supernatant was removed by aspiration, pellets were air dried for 10 min, and then solubilized in 100 µl Tris-HCl (10 mM, pH 8.0). The solutions were pooled, combined with RNase A (2 µl of 20 mg/mL) and incubated at 37 °C for 10 min.

Proteinase K (15 µl) was added to a 1.5 mL tube before the addition of the resuspended precipitated DNA (200 µl) and Buffer AL (200 µl) from the QIAGEN Fast DNA Stool Kit. The mixture was vortexed and incubated at 70 °C for 10 min. Ethanol (200 µl) was added to the mixture and the entire mixture (600 µl) was added to a QIAmp spin column, centrifuged at 16,100 g for 1.5 min, and the flow-through was discarded. The column was then washed with Buffer AW1 (600 µl), centrifuged at 16,100 g for 1.5 min, and the flow-through was discarded. The column was then washed twice with Buffer AW2 (600 µl) as above. The column was centrifuged at 16,100 g for 3 min to remove residual buffer. Pre-heated (65°C) Tris-HCl (100 µl, 10 mM, pH 8.0) was applied to the column, incubated for 1 min, and centrifuged at 16,100 g for 1.5 min to elute the DNA. The yield and purity of DNA was assessed by Nanodrop. DNA was stored at -20°C.

### Blocking efficiency assay

The workflow started with 10 µg of isolated DNA from human fecal material diluted in 30.5 µl filtered H_2_O. As shown in Fig. S1A, one blocking cycle consisted of the following: (**a)** DNA was heated at 94°C for 2 min to denature and immediately cooled on ice for 2 min to disrupt complete re-alignment. (**b)** Heat labile Shrimp Alkaline Phosphatase (rSAP) (1 µl) (NEB #M0371S) was used to remove terminal phosphates that could interfere with terminal extension by incubating in rCutSmart buffer (3.5 µl) (NEB #B6004S) for 30 min at 37°C, and the phosphatase was heat deactivated for 10 min at 65°C. (**c)** Three of four ddNTPs (1 µl, 2 mM each), TriLink BioTechnologies #N- 4001, N-4005, and N-4004) were used to cap available 3’-terminal ends using terminal transferase (1 µl, 20 units) (NEB #M0315S), TdT reaction buffer (1.5 µl), CoCl_2_ (5 µl, 0.25 mM) (NEB #B0252), and filtered H_2_O (4.5 µl) for 60 min at 37°C. We performed this cycle one to seven times on sequential aliquots.

Fresh reagents were added to each subsequent cycle at specific steps: rSAP (1 µl) and rCutSmart buffer (0.3 µl) were added at step b, while ddNTPs (0.3 µl each), Tdt reaction buffer (0.3 µl), CoCl_2_ (0.3 µl), and terminal transferase (1 µl) were added at **Step c**. (**Fig. 1**). After all cycles were complete, excess unincorporated ddNTPs were removed by treatment with a DNA Clean & Concentrator kit (Zymo #11-304C). Finally, any remaining 3’-terminal ends were capped using ddGTP (2mM, TriLink BioTechnologies #N-4002). Excess unincorporated ddGTPs were removed by a DNA Clean & Concentrator kit (Zymo #11- 304C). DNA was then digested as previously published (15) and as outlined below. Incorporated dideoxy components were quantified by LC-MS/MS as outlined below.

### Probe sonication assay

*E. coli* B7A gDNA (10 µg) was diluted in filtered H_2_O (500 µl) (Thermo-Barnstead MicroPure) in a 2 ml Eppendorf tube and kept in an ice-bath at all times to dissipate heat during sonication. DNA was fragmented by probe sonication (Branson Sonifier 250 Ultrasonic Cell Disruptor outfitted with a microtip probe) (50% duty-cycle and an output control setting to attain ∼20-25 on the meter) for 5 cycles (1 cycle = 2 min ON followed by 1 min OFF) (**Fig. S2A**). Fragmented DNA was visualized using 1% agarose gel.

### Iodine cleavage assays

DNA oligos used in this study are listed in **Supplementary Table S1**. 5-mer oligos with a single PT site were ordered from Integrated DNA Technologies (IDT). The 2-mer and 3-mer oligos were ordered from GenScript USA, Inc. The standard curve for each component (2-mer, 3-mer and 5-mer oligos) were calculated based on Beer-Lambert Law @A260 (**Fig. S3**). Separately, retention times for each component were confirmed using MS (Agilent 6490) with no iodine treatment. Reactions (20 µl, 200 pmol oligo, 500 mM Tris-HCl buffer (pH 10.0), 5 mM iodine (Fluka #318981-100), 5 min at 25 °C) were started after adding iodine dilutions which were prepared fresh and used within 2 hrs. We analyzed components, such as the uncut PT oligos and the four 3-mer release oligos (CCA, CCC, CCG, and CCT) using an HPLC system (Agilent 1290) as outlined below, and not by ESI mass spectrometry. Canonical deoxyribonucleosides were also monitored as a normalizing factor.

### Digestion of DNA for LC-MS/MS analysis

Nuclease P1 (United States Biological #N7000) and Calf Intestinal Alkaline Phosphatase (CIAP) (Sigma-Aldrich #P5521) were reconstituted according to the manufacturer’s protocol. DNA (78 µl) was incubated with Nuclease P1 (1.5 U) in 30 mM ammonium acetate pH 5.3 and 0.5 mM ZnCl_2_ (90 µl total) at 55 °C for 2 h. The reaction mixture was then diluted with Tris-HCl (100 mM final concentration, pH 8.0) and incubated with CIAP (51 U) for 2 h at 37 °C. Enzymes were removed by using a VWR 10 kDa spin filter with centrifugation at 12,000 g for 12 min. The solution was dried using a SpeedVac and pellets resuspended in filtered H_2_O (50 µl).

### HPLC analysis of oligos and LC-MS/MS analysis of (dideoxy-) nucleosides and PT dinucleotides

DNA components were analyzed on an Agilent 1290 series HPLC system equipped with a Synergi Fusion RP column (2.5 μm particle size, 100 Å pore size, 100 mm length, 2 mm i.d.) and a diode array detector. The column was eluted at 0.35 mL/min at 35°C with a linear gradient from 3-9% solvent B (acetonitrile) and solvent A (5 mM ammonium acetate, pH 5.3) over 15 min. The column was washed with 95% solvent B for 1 min, and then re-equilibrated with 3% solvent B for 3 min. Oligos and canonical deoxyribonucleosides were quantified by their UV absorbance (A260) using a DAD. In cases where dideoxy- nucleosides and PT dinucleotides were being identified and quantified, the HPLC was coupled to an Agilent 6490 tandem quadrupole mass spectrometry with electrospray ionization operated with the following parameters: N_2_ temperature, 200°C; N_2_ flow rate, 14 L/min; nebulizer pressure, 20 p.s.i.; capillary voltage, 1800 V; and fragmentor voltage, 380 V. For product identification, the mass spectrometer was operated in positive ion multiple reaction monitoring mode using the conditions tabulated in **Supplementary Table S2**.

### Shotgun metagenomic sequencing

We sent an aliquot of human fecal DNA (5 µg) to Novogene Corporation Inc (Sacramento, CA) for shotgun metagenomics sequencing. Briefly, DNA was fragmented and subjected to end repair and phosphorylation after a quality control step. Next, dA-tailing and adaptor ligation was performed using a PCR-free method. Finally, paired-end sequencing (150-bp) was performed on an Illumina Novaseq 6000.

### PTseq.1 library preparation

PTseq.1 was adapted from a previously described protocol (24) (**Fig. 1**). Extracted human fecal DNA (10 µg) was diluted in 500 µl of filtered H_2_O in a 2 ml Eppendorf tube on ice and fragmented by probe sonication (see section above for details). Samples were concentrated using a SpeedVac for ∼2.5 h at ∼450 mTorr and then adjusted with filtered H_2_O to a final volume of 44 µl. Pre-existing strand-breaks were capped with only 1 blocking cycle as outlined above. Briefly, the reaction (50 µl total) contained 10 µg of fragmented DNA, rCutSmart buffer (5 µl), and rSAP (1 µl). Reactions were incubated for 30 min at 37°C to remove 3’-terminal phosphates. The phosphatase was then heat inactivated for 10 min at 65°C. The sample was then filtered using a QIAquick PCR Purification Kit (Qiagen #28106) and eluted with 35 µl elution buffer 10mM Tris-HCl, pH 8.0). Next, the sample was heated to 94 °C in a preheated, hot-lid thermal cycler for exactly 2 min, transferred immediately to ice for 2 min, and then centrifuged briefly. Thereafter, the reaction was adjusted with Tdt reaction buffer (5 µl), CoCl_2_ (5 µl, 0.25 mM), all 4 ddNTPs (1 µl, 2 mM each), and terminal transferase (1 µl, 20 units). The reaction was incubated for 60 min at 37 °C to block any pre-existing strand-breaks. The capped DNA was washed using a DNA Clean & Concentrator kit (Zymo #11-304C).

For subsequent iodine cleavage, the blocked DNA (32 µl) was incubated with 500 mM Tris-HCl pH 9.0 (4 µl) and iodine solution (4 µl, 5 mM) (Fluka, #318981-100) for 10 min at 65 °C and cooled to 4 °C at a speed of 0.1°C/second. The reaction was then split into two 20 µl aliquots and each filtered using a single DyeEX column (QIAGEN #63206) to remove salts and iodine, followed by recombining eluents.

Filtered DNA was again denatured by heating at 94 °C for 2 min and then cooling on ice for 2 min. Thereafter, the reaction was adjusted with rCutSmart buffer (5 µl), rSAP (1 µl), and filtered H_2_O to a final volume of 50 µl, and incubated for 30 min at 37 °C to remove 3ʹ- terminal phosphates arising from iodine cleavage. The phosphatase enzyme was then inactivated for 10 min at 65 °C. The reaction was further adjusted with dTTP (1 µl, 1 mM) (NEB #N0443S), TdT reaction buffer (1 µl), CoCl_2_ (6 µl, 0.25 mM), filtered H_2_O (1 µl), and terminal transferase (1 µl, 20 units) and incubated for 45 min at 37 °C to add dT-tails to the 3’-terminal ends created specifically by iodine at PT sites. The reaction was split into three 20 µl aliquots, excess dTTP was removed using three DyeEX columns, and then eluents were recombined.

The processed DNA was ready for Illumina library preparation using a SMART ChIP-seq kit (Takara, #634865) following the manufacturer’s protocol (Fig. S4). The final PCR step was performed using Illumina primers provided in the kit, and 12 cycles were run for amplification. The PCR product, with unique sequencing barcodes, was submitted to the MIT BioMicro Center for next generation sequencing. Paired-end sequencing (150-bp) was performed on an Illumina MiSeq instrument, using a Custom Read2 sequencing primer that was supplied in the kit (poly(dA) followed by universal read 2 primer).

### PTseq.2 library preparation

PTseq.2 was adapted from PTseq.1 (Fig. 1B) with one adjustment. Human fecal DNA (10 µg) was diluted to 30.5 µl in filtered H_2_O and subjected to four blocking cycles instead of only one blocking cycle. Again, one blocking cycle consisted of the following: **a)** DNA was heated at 94 °C for 2 min to denature and immediately cooled on ice for 2 min. **b)** rSAP (1 µl) and rCutSmart buffer (3.5 µl) were added, the reaction was incubated for 30 min at 37 °C, and the phosphatase was heat deactivated for 10 min at 65 °C **c)** Four ddNTPs (1 µl, 2 mM each), TdT reaction buffer (1.5 µl), CoCl_2_ (5 µl, 0.25 mM), terminal transferase (1 µl, 20 units), and filtered H2O (3.5 µl) were added, and the reaction incubated for 60 min at 37 °C.

Fresh reagents were added to each subsequent cycle at specific steps: rSAP (1 µl) and rCutSmart buffer (0.3 µl) were added at step b, while ddNTPs (0.3 µl each), Tdt reaction buffer (0.3 µl), CoCl_2_ (0.3 µl), and terminal transferase (1 µl) were added at **Step C** (**Fig. 1**). After all cycles were complete, excess unincorporated ddNTPs were removed by treatment with a DNA Clean & Concentrator kit (Zymo #11-304C).

### PTseq.3 library preparation

PTseq.3 was adapted from PTseq.2 (Fig. 1C). Human fecal DNA (10 µg) was diluted in filtered H_2_O and subjected to 4 cycles of blocking as described in PTseq.2. Blocked DNA was then cleaved by iodine at PT sites, washed, and dT-tails were added as described in PTseq.1. Thereafter, the product was diluted in filtered H_2_O (500 µl) and fragmented by probe sonication (as described in the section above). The sample was then concentrated using a SpeedVac to ∼30 µl, followed by Illumina library preparation using a SMART ChIP-seq kit as described in PTseq.1.

### PTseq.4a library preparation

PTseq.4a was adapted from PTseq.3 (Fig. 1D), where all steps were followed through the dT-tailing procedure, including cleaning with three DyeEx columns, as described above. The reaction (60 µl) was then adjusted with TdT reaction buffer (7.8 µl), CoCl_2_ (7.8 µl), ddUTP-biotin (1 µl, 1 mM) (Jena Bioscience #NU-1619-BIOX-S), filtered H_2_O (0.4 µl), and terminal transferase (1 µl) and incubated for 60 min at 37 °C to terminate the dT-tails with a conjugated biotin moiety. After cleaning with four DyeEX columns, and recombining eluents, the processed DNA was diluted in filtered H_2_O (500 µl) and fragmented by probe sonication as described above. Thereafter, DNA fragments were mixed with Hydrophilic Streptavidin Magnetic Beads (5 µl) (NEB #S1421S) and binding buffer (500 µl) (5 mM Tris-HCl, pH 7.5, 1 M NaCl, 0.5 mM EDTA) and incubated for 60 min on a Nutator at room temperature. The beads were subjected to a pull-down process using a magnetic separation rack (Dynal #MPC-S) and washed three times with binding buffer (100 µl). After discarding the supernatant, beads were resuspended in filtered H_2_O (20 µl).

Finally, the enriched DNA that was captured on the beads was directly subjected to Illumina library preparation using a SMART ChIP-seq kit as described in PTseq.1.

### PTseq.4b library preparation

PTseq.4b was adapted from PTseq.4a, where differences only emerged after bead capture (**Fig. 1D Step E**, **Fig. S4**). After washing, the captured beads were resuspended in a reaction with oligo (dA) primer (5 µl, 50 pmol), and filtered H_2_O (20 µl). The reaction was heated for 1 min at 94 °C and immediately transferred to ice for exactly 2 min. The reaction was then adjusted with PrimeScript buffer (10 µl), dNTP mixture (5 µl, 1 mM each), PrimeScript reverse transcriptase (4 µl) (Takara, #2680B), and filtered water (6 µl) to a final volume of 50 µl to synthesize the first cDNA strand. The reaction was incubated for 60 min at 42 °C. The reaction was then heated for 15 min at 70 °C to deactivate the reverse transcriptase, and cooled on ice. Finally, the product was sent to Novogene Corporation, Inc (Sacramento, CA) where standard, ligation-based library preparation was performed using a NEBNext Ultra II DNA Kit (NEB). The library was then subjected to Illumina NovaSeq paired-end sequencing (150-bp).

### PT-seq data analysis of bacterial gDNA

**Fig. S5A** provides a graphical overview of the data analysis process. We used a series of bioinformatic tools included with the BBMap package at sourceforge.net. For example, dT-tails and adapters (Read1: AGATCGGAAGAGCGTCGTGTAGGGAAAGAGTGT and Read2: TTTTTTTTTTTTTTTAGATCGGAAGAGCACACGTCTGAACTCCAGTCA) were removed using the BBDuk tool with the following parameters: ktrim=r k=18 hdist=2 hdist2=1 rcomp=f mink=8 qtrim=r trimq=30 for R1, ktrim=r k=18 mink=8 hdist=1 rcomp=f qtrim=r trimq=30 for R2. PhiX was removed using BBDuk with parameters k=31 hdist=1. Trimmed reads were aligned to genome sequences using Bowtie 2 with parameters set to: sensitive. The genome sequence of *Bacteroides salyersiae* DSM 18765 was downloaded from NCBI RefSeq (GCF_000381365.1). The genome sequences of *Lachnospiraceae sp.* and *Butyricimonas faecalis* were provided by the GMbC collection (33). The bam files were cleaned using SMARTcleaner (34) and divided into two strands using samtools with view settings -f 163 and -f 147, respectively (35). The coverage of each strand was calculated separately using bedtools (36) with the following setting: genomecov -d -5. Thereafter, custom scripts were used for pileup calling (scripts are available upon request). Briefly, the start positions of all individual reads were recorded for each strand separately. The read-pileup depth at each position was represented by the number of reads starting at each position (from the 5’-end). The read-pileups for which the depth was more than half of the sequencing depth were retained. The 13 nt sequences centered at positions with depth more than 1 were retrieved using samtools faidx (35). The consensus motifs were analyzed with incremental depth, typically from 1 to 200 (stepwise by 10), using MEME (37) with parameters -dna - objfun classic -nmotifs 5 -mod zoops -evt 0.05 -minw 3 -maxw 4 -markov_order 0 -nostatus - oc. Pileups within < 3 bps were collapsed into the centermost consensus motif. The read-pileup depth cutoff was determined to generate pileups that were equivalent to the level of PTs that were quantified by LC-MS/MS analysis. Specificity was calculated by the percentage of pileups at the conserved motif sites among all pileup sites.

### Data analysis of PT-seq of human fecal DNA

The steps for the generation of reference genomes are indicated by blue arrows in **Fig. S5B**. First, bacterial and archaeal genomes were retrieved from the Broad Institute-OpenBiome Microbiome Library (BIO-ML) (38), the Global Microbiome Conservancy (GMbC) collection (33), and the Unified Human Gastrointestinal Genome (UHGG) collection, respectively (39). The 13,663 genomes were processed by dRep to remove duplications (40), resulting in a set of 5,067 best representative genomes. The reads from shotgun metagenomic sequencing (see above section) were then assigned to the 5,067 genomes using Kraken2 (41) and the abundance was estimated using Bracken (42). The top 200 most abundant genomes were used as reference genomes for mapping PTseq (**Supplementary Table S3**).

**Fig. S5B** (black arrows) provides a graphical overview of analyzing PTseq reads. Reads were trimmed as described above. Trimmed read pairs were aligned to reference genomes using Mapper v1.1.0-beta05 (https://github.com/mathjeff/Mapper). If a specific read maps to multiple positions (assuming optimal alignment quality), we output all alignment results and calculated depth to each position as 1/n (n=number of optimal aligned positions). The start positions of Read2 reads that represent iodine cleaved sites were located by comparison to the alignment with only Read2 reads. The resulting alignments in vcf format were converted to locations of read-pileup in tab delimited text format using custom scripts (scripts are available upon request). Then, a sequence window (17-nt) centered at positions with depth more than 1 were retrieved and conserved motifs were analyzed as described above. Genomes with conserved motifs detected were considered as PT-containing genomes. Finally, a read-pileup depth cutoff was determined in PT-containing genomes to generate pileups that were equivalent to the level of PTs quantified by LC-MS/MS analysis (note: the number of pileups for each genome was weighted by genome length and abundance).

### Full-length 16S amplicon sequencing

One aliquot of human fecal DNA (2 µg) was cleaved by iodine as described in PTseq.1 above (**Fig. S6**). A control aliquot of DNA was treated with water instead of iodine. After washing, samples were directly submitted to a 16S Amplicon sequencing service at Zymo Research (Irvine, CA). Briefly, a library was prepared for each sample by following the full-length 16S amplification protocol outlined by Pacific Bioscience (Menlo Park, CA). The whole 16S gene was amplified using 27f (AGRGTTYGATYMTGGCTCAG) and 1492r (RGYTACCTTGTTACGACTT) primers with barcodes and adapters. After PCR amplification and quality control, samples were pooled and processed using a SMRTbell prep kit 3.0 (Pacific Bioscience, Menlo Park, CA). Finally, the library was sequenced using an 8M SMRT cell on a Sequal IIe system for 15 hours. Bioinformatic analyses were performed by Zymo Research with QIIME v.1.9.1 (43).

## DATA AVAILABILITY

The custom scripts for processing the sequencing data as described in Methods are available upon request. Sequencing data has been deposited in NCBI SRA database under BioProject ID PRJNA1006039.

## SUPPLEMENTARY DATA

Supplementary data are available online.

## AUTHOR CONTRIBUTIONS

Yifeng Yuan and Michael S. DeMott: conceptualized, designed, and performed the experiments, completed data analysis, interpreted results, and wrote the manuscript. Shane R. Byrne: assisted with experimental analysis and method development. Peter C. Dedon: Conceived of the project, supervised all aspects of this process, and reviewed, edited, and finalized the manuscript.

## ACKNOWLEDGEMENTS

We thank Dr. Laurie Comstock (University of Chicago, Chicago, IL) for providing bacterial cell pellets. We thank Susan Weir and Katya Moniz at Openbiome for assisting with bacterial isolates. We thank the MIT BioMicro Center and its Director, Dr. Stuart Levine, for support and advice during the performance of the studies presented here.

## REFERENCES

1. Sanchez-Romero, M.A. and Casadesus, J. (2020) The bacterial epigenome. Nat Rev Microbiol, 18, 7–20.

2. Weigele, P. and Raleigh, E.A. (2016) Biosynthesis and Function of Modified Bases in Bacteria and Their Viruses. Chem Rev, 116, 12655–12687.

3. Gish, G. and Eckstein, F. (1988) DNA and RNA sequence determination based on phosphorothioate chemistry. Science, 240, 1520–1522.

4. Xie, X., Liang, J., Pu, T., Xu, F., Yao, F., Yang, Y., Zhao, Y.L., You, D., Zhou, X., Deng, Z. et al. (2012) Phosphorothioate DNA as an antioxidant in bacteria. Nucleic Acids Res, 40, 9115–9124.

5. Cao, B., Chen, C., DeMott, M.S., Cheng, Q., Clark, T.A., Xiong, X., Zheng, X., Butty, V., Levine, S.S., Yuan, G. et al. (2014) Genomic mapping of phosphorothioates reveals partial modification of short consensus sequences. Nat Commun, 5, 3951.

6. Dai, D., Du, A., Xiong, K., Pu, T., Zhou, X., Deng, Z., Liang, J., He, X. and Wang, Z. (2016) DNA Phosphorothioate Modification Plays a Role in Peroxides Resistance in Streptomyces lividans. Front Microbiol, 7, 1380.

7. Kellner, S., DeMott, M.S., Cheng, C.P., Russell, B.S., Cao, B., You, D. and Dedon, P.C. (2017) Oxidation of phosphorothioate DNA modifications leads to lethal genomic instability. Nat Chem Biol, 13, 888–894.

8. Tong, T., Chen, S., Wang, L., Tang, Y., Ryu, J.Y., Jiang, S., Wu, X., Chen, C., Luo, J. and Deng, Z. (2018) Occurrence, evolution, and functions of DNA phosphorothioate epigenetics in bacteria. Proceedings of the National Academy of Sciences, 115, E2988–E2996.

9. Li, J., Chen, Y., Zheng, T., Kong, L., Zhu, S., Sun, Y., Deng, Z., Yang, L. and You, D. (2019) Quantitative mapping of DNA phosphorothioatome reveals phosphorothioate heterogeneity of low modification frequency. PLoS Genet, 15, e1008026.

10. Wang, L., Jiang, S., Deng, Z., Dedon, P.C. and Chen, S. (2019) DNA phosphorothioate modification—a new multi-functional epigenetic system in bacteria. FEMS microbiology reviews, 43, 109–122.

11. Sun, Y., Kong, L., Wu, G., Cao, B., Pang, X., Deng, Z., Dedon, P.C., Zhang, C. and You, D. (2020) DNA Phosphorothioate Modifications Are Widely Distributed in the Human Microbiome. Biomolecules, 10.

12. Zhu, S., Zheng, T., Kong, L., Li, J., Cao, B., DeMott, M.S., Sun, Y., Chen, Y., Deng, Z., Dedon, P.C. et al. (2020) Development of Methods Derived from Iodine-Induced Specific Cleavage for Identification and Quantitation of DNA Phosphorothioate Modifications. Biomolecules, 10.

13. Wang, S., Wan, M., Huang, R., Zhang, Y., Xie, Y., Wei, Y., Ahmad, M., Wu, D., Hong, Y., Deng, Z. et al. (2021) SspABCD-SspFGH Constitutes a New Type of DNA Phosphorothioate-Based Bacterial Defense System. mBio, 12.

14. Huang, Q., Lee, G.Y., Li, J., Wang, C., Zhao, R.L., Deng, Z., Houk, K.N. and Zhao, Y.L. (2022) Origin of iodine preferential attack at sulfur in phosphorothioate and subsequent P-O or P-S bond dissociation. Proc Natl Acad Sci U S A, 119, e2119032119.

15. Byrne, S., DeMott, M., Yuan, Y., Ghanegolmohammadi, F., Kaiser, S., Fox, J., Alm, E. and Dedon, P. (2024) Temporal dynamics and metagenomics of phosphorothioate epigenomes in the human gut microbiome bioRxiv.

16. Yuan, Y., DeMott, M., Byrne, S., Flores, K., Poyet, M., Berdy, B., Comstock, L., Alm, E. and Dedon, P. (2024) Human gut microbiome comparative genomics reveals new phosphorothioate epigenetics systems. bioRxiv.

17. Wang, L., Chen, S., Vergin, K.L., Giovannoni, S.J., Chan, S.W., DeMott, M.S., Taghizadeh, K., Cordero, O.X., Cutler, M., Timberlake, S. et al. (2011) DNA phosphorothioation is widespread and quantized in bacterial genomes. Proc Natl Acad Sci U S A, 108, 2963–2968.

18. Wang, L., Chen, S., Xu, T., Taghizadeh, K., Wishnok, J.S., Zhou, X., You, D., Deng, Z. and Dedon, P.C. (2007) Phosphorothioation of DNA in bacteria by dnd genes. Nat Chem Biol, 3, 709–710.

19. Wang, L., Jiang, S., Deng, Z., Dedon, P.C. and Chen, S. (2019) DNA phosphorothioate modification-a new multi-functional epigenetic system in bacteria. FEMS Microbiol Rev, 43, 109–122.

20. Xiong, L., Liu, S., Chen, S., Xiao, Y., Zhu, B., Gao, Y., Zhang, Y., Chen, B., Luo, J., Deng, Z. et al. (2019) A new type of DNA phosphorothioation-based antiviral system in archaea. Nat Commun, 10, 1688.

21. Huang, Q., Li, J., Shi, T., Liang, J., Wang, Z., Bai, L., Deng, Z. and Zhao, Y.L. (2020) Defense Mechanism of Phosphorothioated DNA under Peroxynitrite-Mediated Oxidative Stress. ACS Chem Biol, 15, 2558–2567.

22. Dedon, P.C. and Tannenbaum, S.R. (2004) Reactive nitrogen species in the chemical biology of inflammation. Arch Biochem Biophys, 423, 12–22.

23. Bhattacharyya, A., Chattopadhyay, R., Mitra, S. and Crowe, S.E. (2014) Oxidative stress: an essential factor in the pathogenesis of gastrointestinal mucosal diseases. Physiol Rev, 94, 329–354.

24. Cao, B., Wu, X., Zhou, J., Wu, H., Liu, L., Zhang, Q., DeMott, M.S., Gu, C., Wang, L., You, D. et al. (2020) Nick-seq for single-nucleotide resolution genomic maps of DNA modifications and damage. Nucleic Acids Res.

25. Jayaprakash, A.D., Jabado, O., Brown, B.D. and Sachidanandam, R. (2011) Identification and remediation of biases in the activity of RNA ligases in small-RNA deep sequencing. Nucleic Acids Res, 39, e141.

26. Cai, W.M., Chionh, Y.H., Hia, F., Gu, C., Kellner, S., McBee, M.E., Ng, C.S., Pang, Y.L., Prestwich, E.G., Lim, K.S. et al. (2015) A Platform for Discovery and Quantification of Modified Ribonucleosides in RNA: Application to Stress-Induced Reprogramming of tRNA Modifications. Methods in enzymology, 560, 29–71.

27. Milowska, K. and Gabryelak, T. (2007) Reactive oxygen species and DNA damage after ultrasound exposure. Biomol Eng, 24, 263–267.

28. Nakamaye, K.L., Gish, G., Eckstein, F. and Vosberg, H.P. (1988) Direct sequencing of polymerase chain reaction amplified DNA fragments through the incorporation of deoxynucleoside alpha-thiotriphosphates. Nucleic Acids Res, 16, 9947–9959.

29. Motea, E.A. and Berdis, A.J. (2010) Terminal deoxynucleotidyl transferase: The story of a misguided DNA polymerase. Biochimica et Biophysica Acta (BBA) - Proteins and Proteomics, 1804, 1151–1166.

30. Yang, J., Pu, J., Lu, S., Bai, X., Wu, Y., Jin, D., Cheng, Y., Zhang, G., Zhu, W., Luo, X., et al. (2020) Species-Level Analysis of Human Gut Microbiota With Metataxonomics. Front Microbiol, 11, 2029.

31. McCallum, G. and Tropini, C. (2024) The gut microbiota and its biogeography. Nat Rev Microbiol, 22, 105–118.

32. Sender, R., Fuchs, S. and Milo, R. (2016) Revised Estimates for the Number of Human and Bacteria Cells in the Body. PLoS Biol, 14, e1002533.

33. Groussin, M., Poyet, M., Sistiaga, A., Kearney, S.M., Moniz, K., Noel, M., Hooker, J., Gibbons, S.M., Segurel, L., Froment, A. et al. (2021) Elevated rates of horizontal gene transfer in the industrialized human microbiome. Cell, 184, 2053–2067 e2018.

34. Zhao, D. and Zheng, D. (2018) SMARTcleaner: identify and clean off-target signals in SMART ChIP-seq analysis. BMC Bioinformatics, 19, 544.

35. Li, H., Handsaker, B., Wysoker, A., Fennell, T., Ruan, J., Homer, N., Marth, G., Abecasis, G., Durbin, R. and Genome Project Data Processing, S. (2009) The Sequence Alignment/Map format and SAMtools. Bioinformatics, 25, 2078–2079.

36. Quinlan, A.R. and Hall, I.M. (2010) BEDTools: a flexible suite of utilities for comparing genomic features. Bioinformatics, 26, 841–842.

37. Bailey, T.L., Johnson, J., Grant, C.E. and Noble, W.S. (2015) The MEME Suite. Nucleic Acids Research, 43, W39–W49.

38. Poyet, M., Groussin, M., Gibbons, S.M., Avila-Pacheco, J., Jiang, X., Kearney, S.M., Perrotta, A.R., Berdy, B., Zhao, S., Lieberman, T.D. et al. (2019) A library of human gut bacterial isolates paired with longitudinal multiomics data enables mechanistic microbiome research. Nat Med, 25, 1442–1452.

39. Almeida, A., Nayfach, S., Boland, M., Strozzi, F., Beracochea, M., Shi, Z.J., Pollard, K.S., Sakharova, E., Parks, D.H., Hugenholtz, P. et al. (2021) A unified catalog of 204,938 reference genomes from the human gut microbiome. Nat Biotechnol, 39, 105–114.

40. Olm, M.R., Brown, C.T., Brooks, B. and Banfield, J.F. (2017) dRep: a tool for fast and accurate genomic comparisons that enables improved genome recovery from metagenomes through de-replication. ISME J, 11, 2864–2868.

41. Lu, J., Rincon, N., Wood, D.E., Breitwieser, F.P., Pockrandt, C., Langmead, B., Salzberg, S.L. and Steinegger, M. (2022) Metagenome analysis using the Kraken software suite. Nat Protoc, 17, 2815–2839.

42. Lu, J., Breitwieser, F.P., Thielen, P. and Salzberg, S.L. (2017) Bracken: estimating species abundance in metagenomics data. PeerJ Computer Science, 3.

43. Caporaso, J.G., Kuczynski, J., Stombaugh, J., Bittinger, K., Bushman, F.D., Costello, E.K., Fierer, N., Pena, A.G., Goodrich, J.K., Gordon, J.I. et al. (2010) QIIME allows analysis of high-throughput community sequencing data. Nat Methods, 7, 335–336.

