## Supplementary Figures for "PT-seq: A method for metagenomic analysis of phosphorothioate epigenetics in complex microbial communities"

**Contents**

Figure S1. Effect of repeated blocking cycles on 3′-ends

Figure S2. Test of probe sonication as method of fragmenting DNA

Figure S3. Scheme for measuring iodine cutting efficiency in DNA oligos

Figure S4. The scheme of library prep of PTseq.4a and PTseq.4b

Figure S5. Data processing workflow for PTseq

Figure S6. Workflow for microbiome full-length 16S amplicon PT-seq

Figure S7. Concentration, sequence, and structure dependence of iodine cleavage at PTs

Figure S8. Optimizing temperature, time, and Tris conditions for the iodine cleavage reaction

Figure S9. The low sensitivity and specificity of PTseq.2 in bacterial isolate genomes

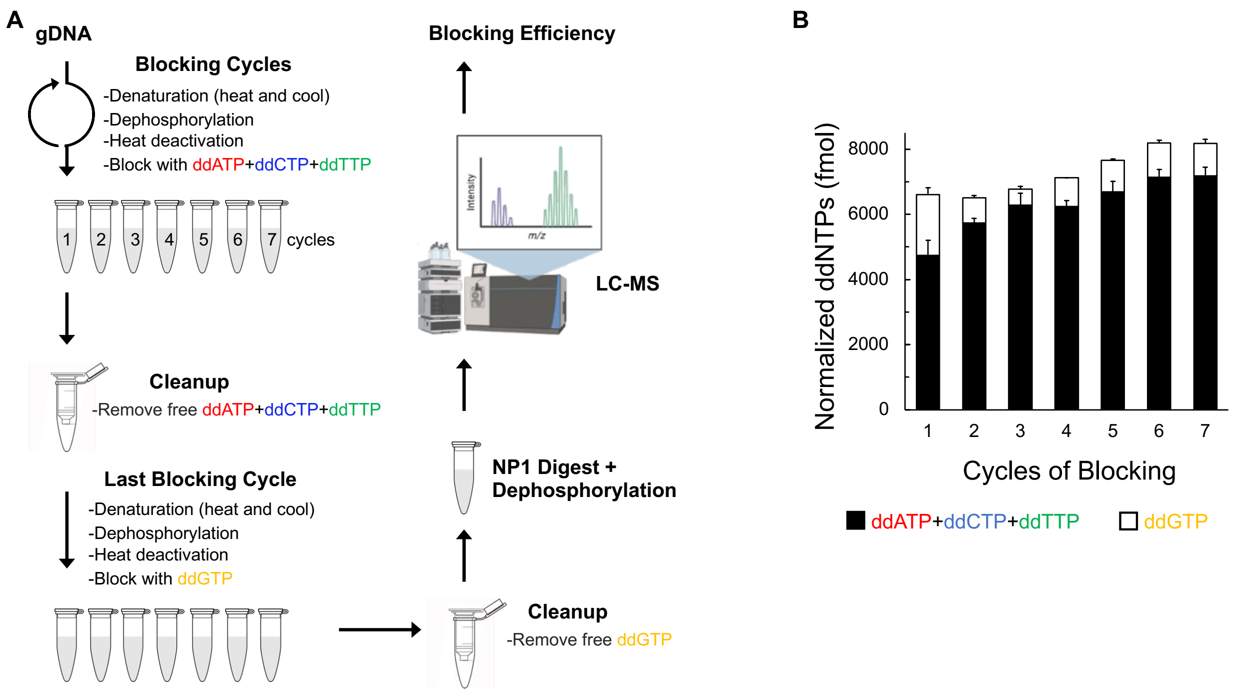


**Fig. S1. Effect of repeated blocking cycles on 3′-ends.** (A) The scheme of measuring the incorporated ddNTPs in 10 µg of human gut microbiome DNA after repeated blocking cycles. One blocking cycle consisted of the following: 1) denaturation of DNA by heating and rapidly cooling on ice, 2) removal of terminal phosphates using a heat labile phosphatase, 3) heat deactivation of enzyme, 4) three of four ddNTPs were used to cap and block available 3′-terminal ends. (B) The number of incorporated ddATP, ddCTP and ddTTP increased with repeated cycles, suggesting that repeated blocking cycles reduce available 3′-OH terminal ends for erroneous downstream T-tailing. The number of ddGTP did not increase, indicating that sample handling did not create new 3′-ends.

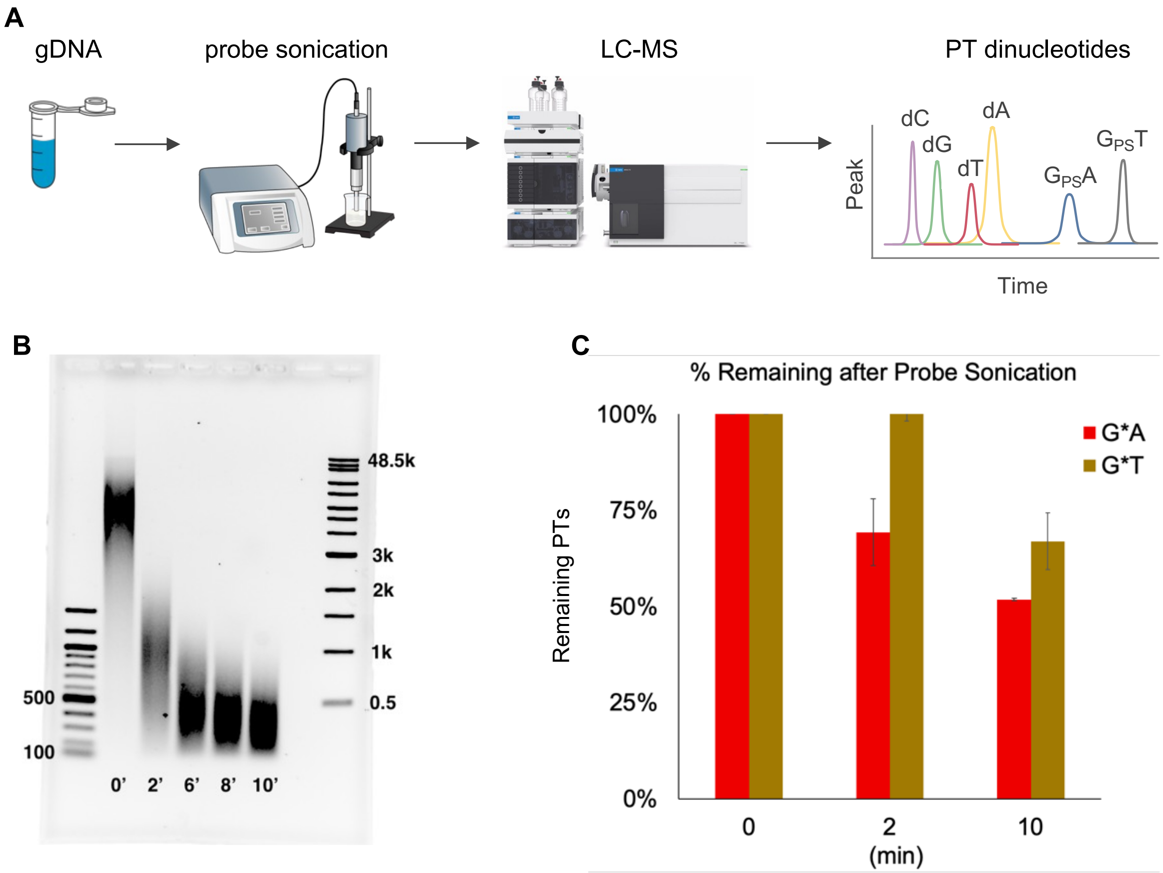


**Fig. S2. Test of probe sonication as method of fragmenting DNA.** (A) Scheme for measuring remaining PTs in the sample. (B) Agarose gel showing probe sonication time course (0-10 min) of human fecal DNA. (C) Bar graphs showing % remaining of PT dinucleotides (G*A) and (G*T) as measured by LC-MS in gDNA from *E. coli* B7A after probe sonication for 0, 2, and 10 min.

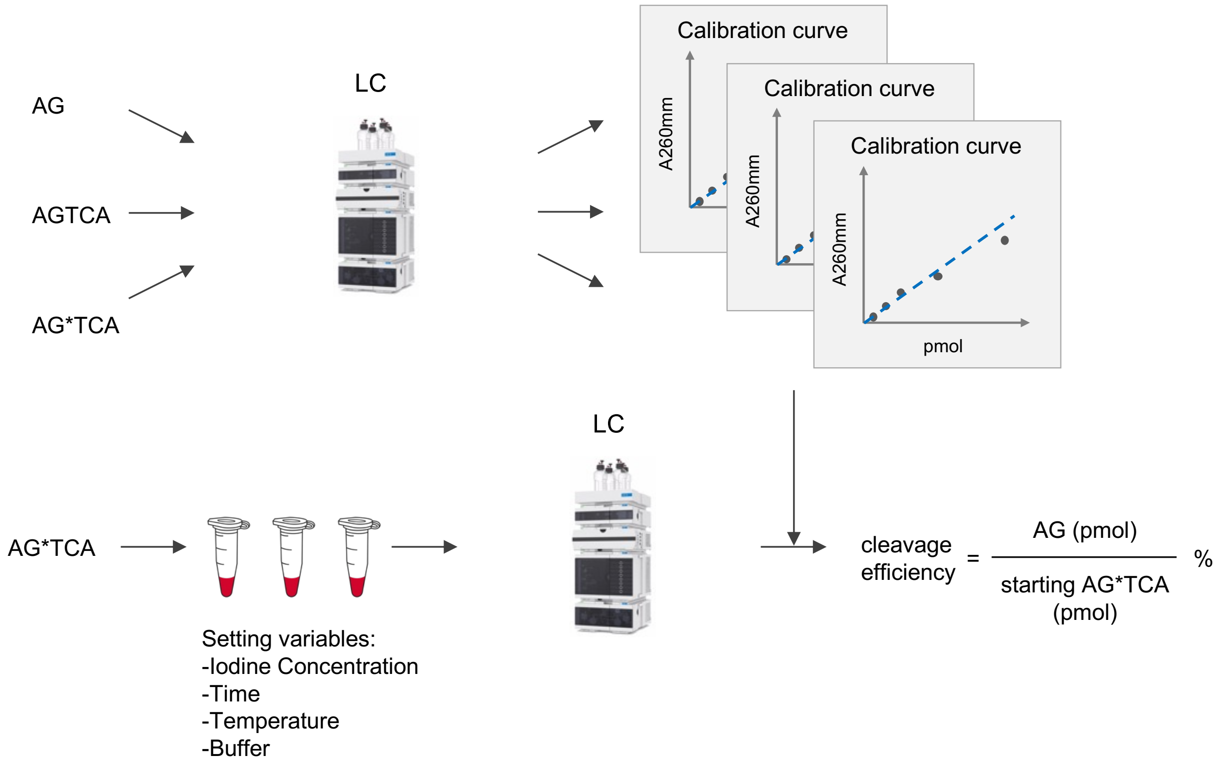


**Fig. S3. Scheme for measuring iodine cutting efficiency in DNA oligos.** The elution of each component using an HPLC system and calibration curves were characterized using UV A260mm. The amount of starting oligos and released product were measured to calculate cutting efficiency.

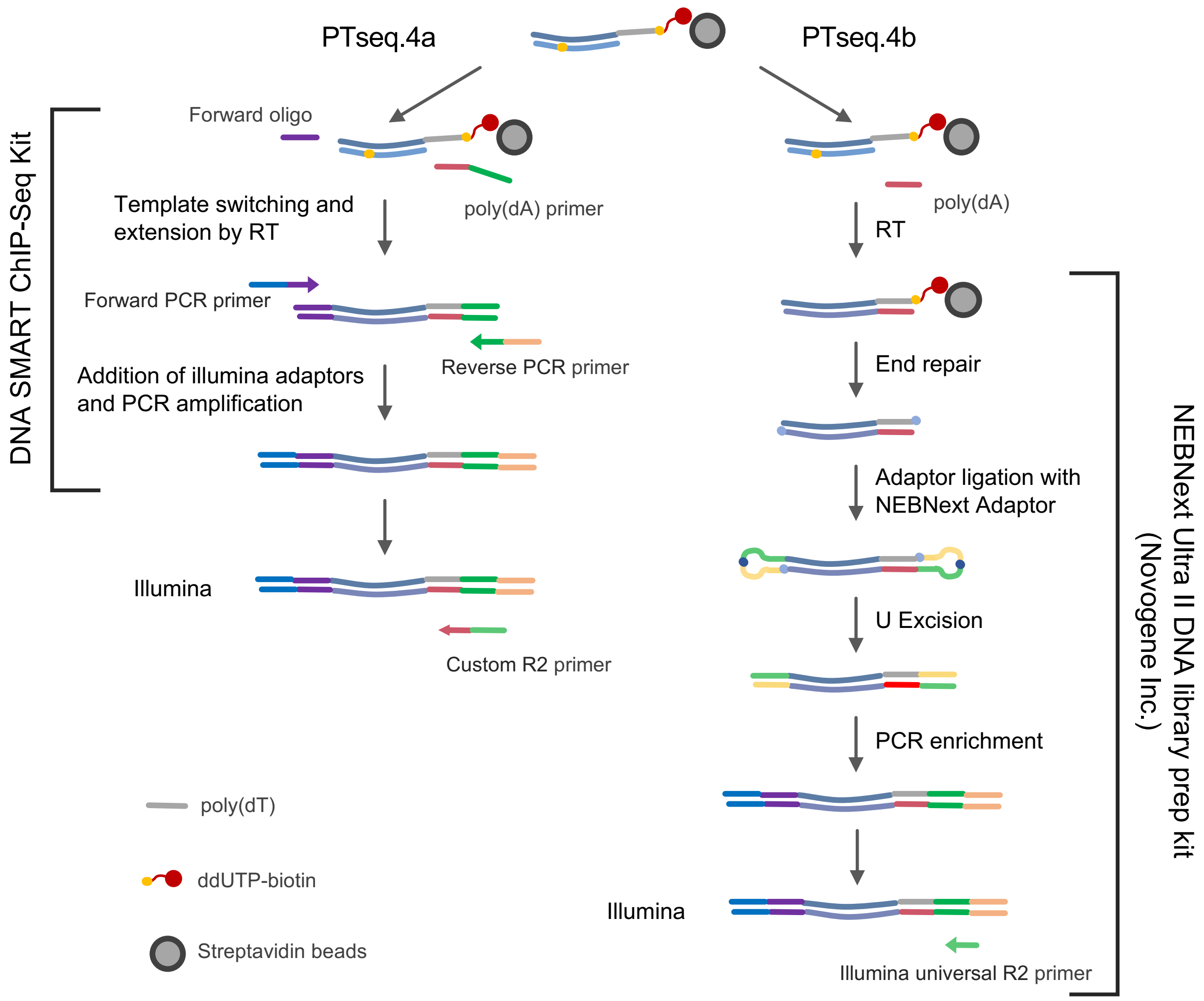


**Fig. S4. The scheme of library prep of PTseq.4a and PTseq.4b.** Library prep of PTseq.4a followed the manual of DNA SMART ChIP-seq kit. Library prep of PTseq.4b was done by Novogene Inc. using NEBNext Ultra II DNA library prep kit.

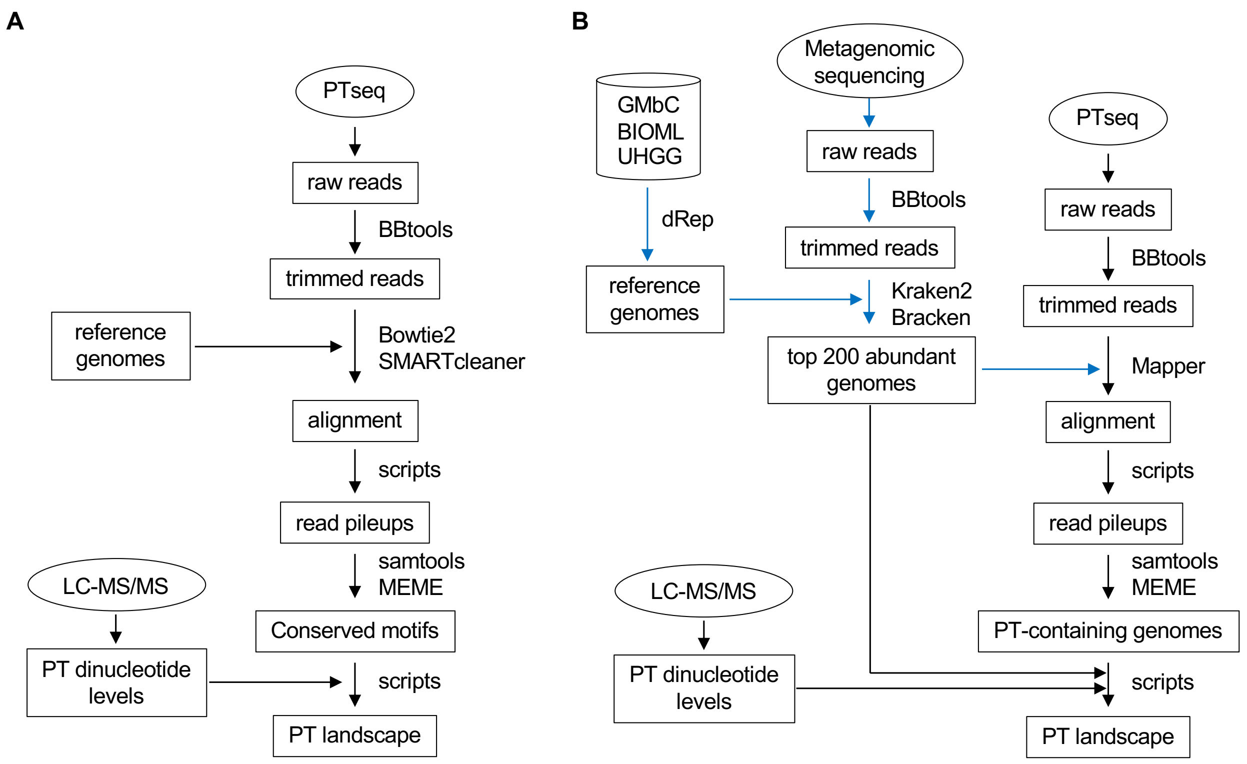


**Fig. S5. Data processing workflow for PTseq.** Blue arrows represent reference genome preparation using metagenomic sequencing data. Black arrows represent data processing of PTseq in bacterial isolates (A) and in human gut microbiome sample (B).

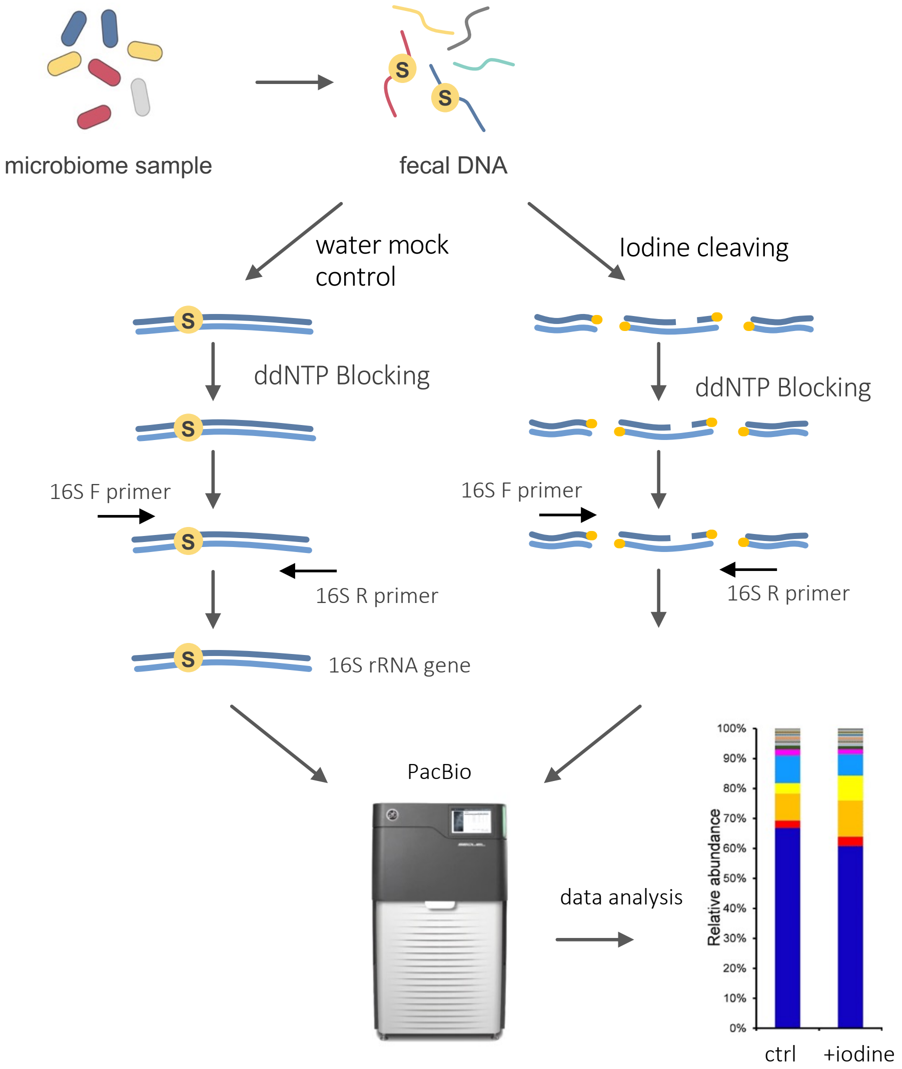


**Fig. S6. Workflow for microbiome full-length 16S amplicon PT-seq.** Human fecal DNA was aliquoted for iodine cleavage and water mock control. Full-length 16S rRNA genes cannot be PCR amplified so removed from library. Comparison of OTU abundance reveals PT-containing 16S rRNA genes.

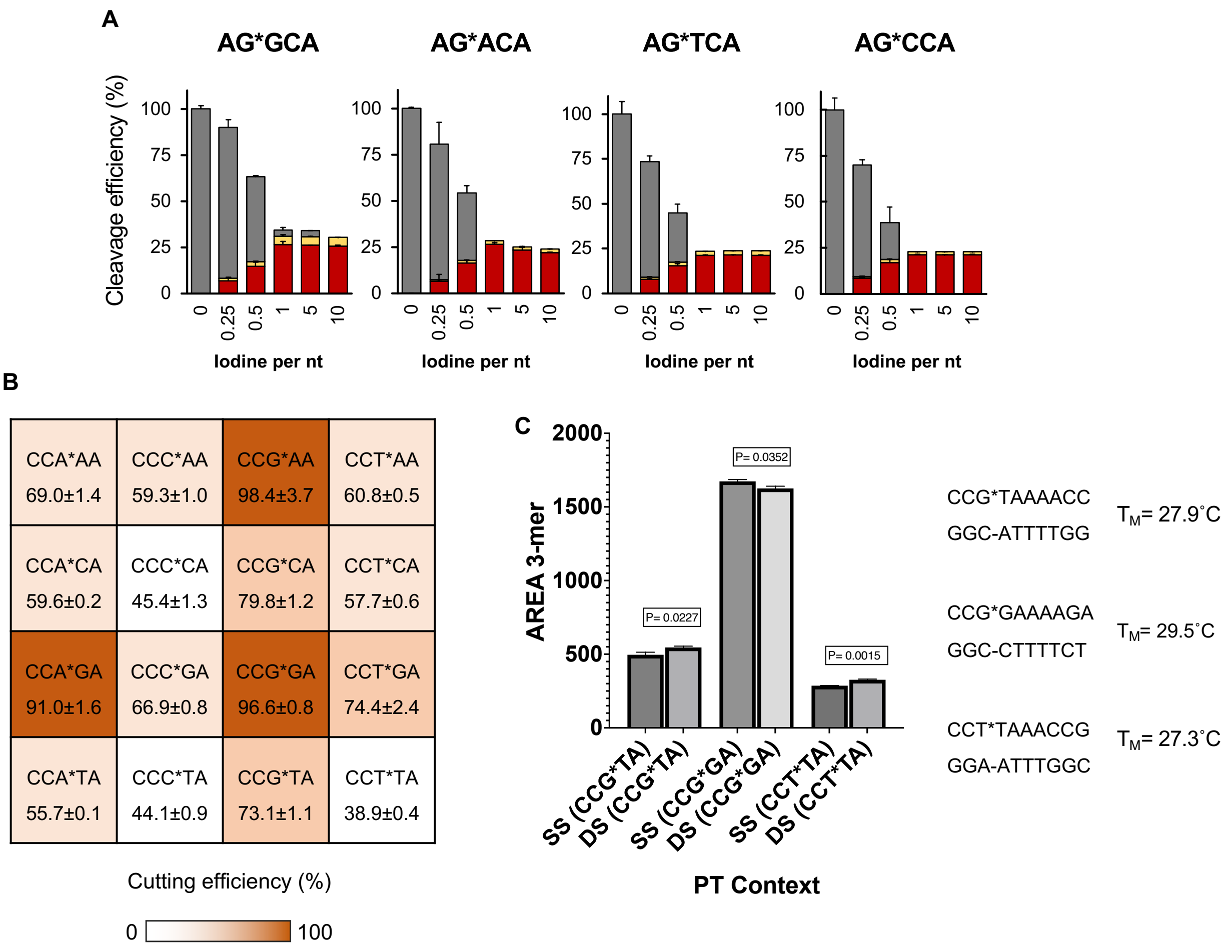


**Fig. S7. Concentration, sequence, and structure dependence of iodine cleavage at PTs.** (**A**) Effect of iodine concentration on cutting efficiency of PTs. 5-mer PT oligos were incubated with increasing iodine concentrations (relative to nucleotides) at room temperature for 5 min. (**B**) Effect of all 16 PT dinucleotide sequence contexts on 5 mM iodine cutting efficiency for single-stranded DNA. (**C**) Comparisons of 5 mM iodine cutting efficiencies for the single- and double-stranded forms of three representative sequence context of low, median, high cutting efficiency. The melting temperatures for the three sequence contexts in double-stranded form. Reactions were performed at 15˚C. Data represent mean ± SD for three replicates.

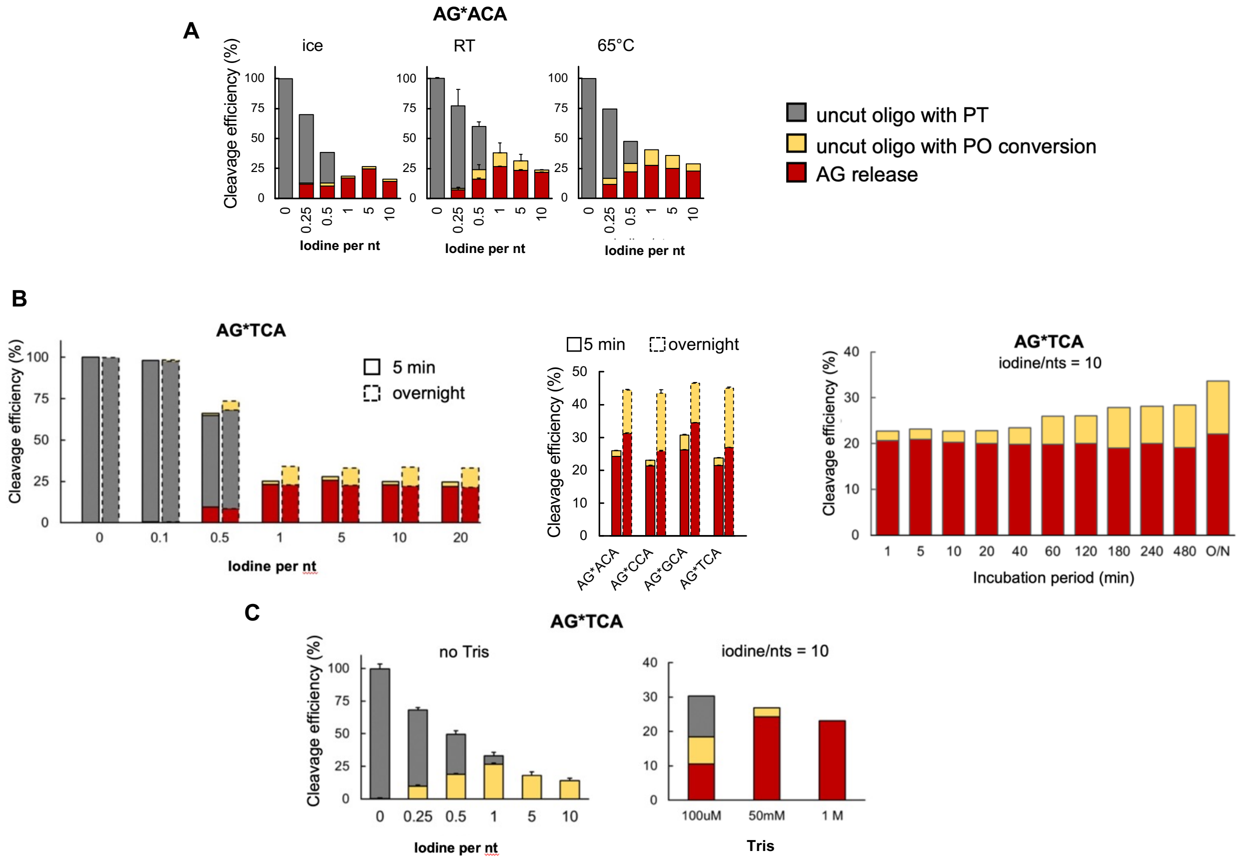


**Fig. S8. Optimizing temperature, time, and Tris conditions for the iodine cleavage reaction.** (**A**) Comparison of incubation temperature (ice, room temperature/RT, 65 ° C) on iodine cutting efficiency of PTs in a 5-min reaction with AG*ACA. (**B**) Effect of incubation time on iodine cutting efficiency at different iodine concentrations with AG*TCA. Left: AG*TCA was incubated at RT for 5 min and overnight with different iodine concentrations. Middle: Four oligos were incubated at RT for 5 min and overnight with an iodine:nt ratio of 10. Right: Time course of iodine cutting efficiency. (**C**) Effect of Tris concentration on iodine cutting efficiency with AG*TCA. Data represent mean±SD for three replicates.

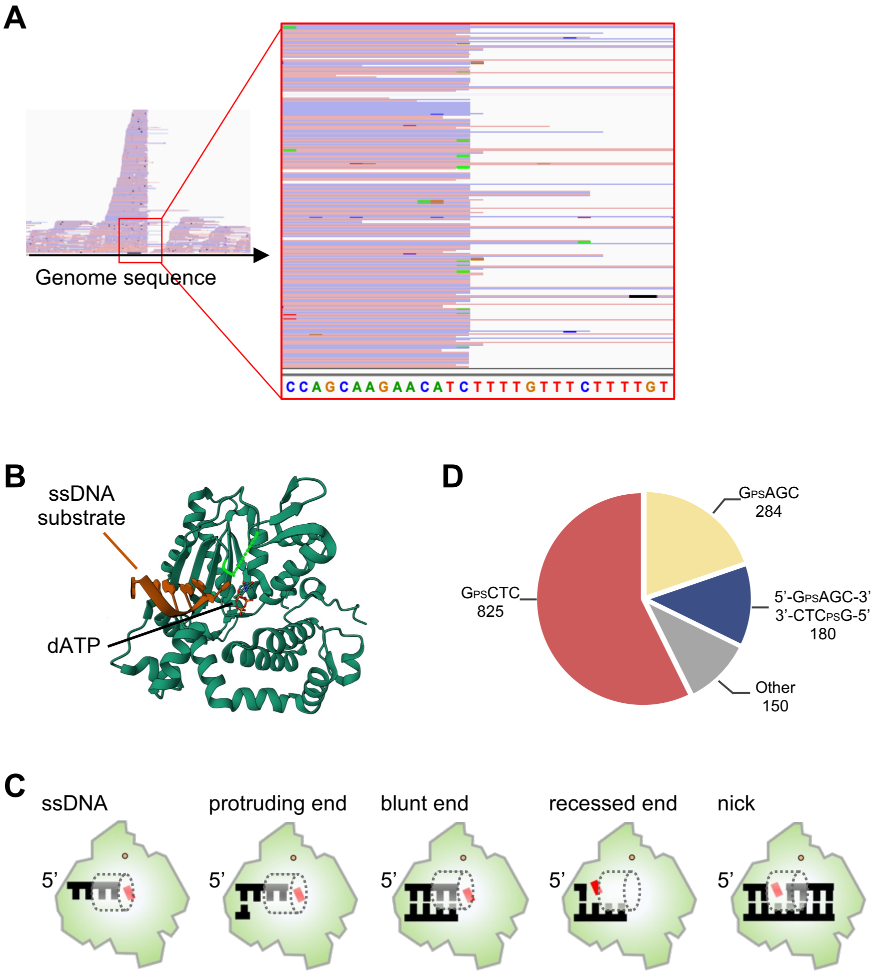


**Fig. S9. The low sensitivity and specificity of PTseq.2 in bacterial isolate genomes.**
(A) An example of read alignment at a polyT site in *Lachnospiraceae sp.* in Integrative Genomics Viewer. (B) The protein structure of mouse terminal transferase (PDB 4I27) in green cartoon. The ssDNA substrate is in brown. dATP is shown in sticks. (C) The schematic illustration of the structure of mouse terminal transferase. Single and double DNA strands are indicated in black. The coming deoxynucleotide is indicated in the red block. The dot indicates an ion. The dashed tube indicates the space inside the lariat-like structure that prevents the ability of TdT to accommodate a templating strand. (D) The number of PT modification sites identified by original ICDS. A total of 1,439 PT sites were detected, with 1,289 of PT sites occurred at either single stranded or double stranded GAGC/GCTC sites.
